# eeeHive: a new HF RFID-based automated behavioral monitoring system for group-housed animals with high spatiotemporal resolution

**DOI:** 10.64898/2026.04.30.720993

**Authors:** Seico Benner, Suzuka Shiono, Toshinori Kagawa, Kiyohiko Hattori, Hidenori Yamasue, Hans-Peter Lipp, Toshihiro Endo

**Affiliations:** Center for Health and Environmental Risk Research, National Institute for Environmental Studies, Tsukuba, Japan; Department of Psychiatry, Hamamatsu University School of Medicine, Hamamatsu, Japan; Phenovance LLC, Kashiwa, Japan; Grid and Communication Technology Division, Grid Innovation Research Laboratory, Central Research Institute of Electric Power Industry, Yokosuka, Japan; Department of Information Systems and Multimedia Design, School of Science and Technology for Future Life, Tokyo Denki University, Tokyo, Japan; Faculty of Medicine, Institute of Evolutionary Medicine, University of Zürich, Zürich, Switzerland

**Keywords:** Automated Home-Cage Monitoring (AHCM)_1_, RFID_2_, high-frequency (HF) RFID_3_, ISO/IEC 15693_4_, eeeHive_5_, Ethological approach_6_, Group-housing_7_, Animal welfare_8_

## Abstract

Long-term, automated tracking of group-housed social animals using RFID (radio frequency identification) is a promising approach in ethological neuroscience. However, low-frequency (LF) RFID, while long-established in the field, is constrained by its inherent low data rates, which lead to two critical limitations: (1) compromised spatiotemporal resolution, and (2) the inability to identify multiple tags (animals) simultaneously. To address these limitations, we developed eeeHive, a high-frequency (HF) RFID-based animal tracking system with a fully custom hardware architecture that enables high-speed, multiplexed antenna polling and concurrent multi-tag reading. The polling time per antenna in eeeHive was 5.9 ms, with an additional 8.2 ms read time per tag. We applied the system to track 24 mice for one week, and six common marmosets for seven weeks. The system successfully tracked individuals even within dense clusters, revealing complex behavioral traits characterized by spatial utilization, temporal dynamics, behavioral regularity, and inter-individual relationships. Additional tests with Japanese fire-bellied newts and Nile tilapia juveniles demonstrated comparable tracking performance in aquatic environments. Taken together, eeeHive overcomes the inherent limitations of conventional LF RFID, establishing a powerful HF RFID-based platform for fine-scale behavioral tracking of group-housed animals across terrestrial and aquatic species.

## 1 Introduction

The diversity and availability of model animals in neuroscience and neuropharmacology have expanded substantially over the past two decades (1,2). However, the reliable interpretation of their behavior in laboratory environments remains a persistent challenge (3–6). Conventional behavioral test paradigms, particularly in rodents, typically involve manually transferring animals from their home cages to isolated testing chambers for short observation periods (7). Since around the 2000s, there has been criticism of such “closed-session” procedures for relying on operational definitions that disregard the impact of human intervention and the sensory and behavioral traits unique to each species or strain (8,9).

To address this problem, the concept of automated home-cage monitoring (AHCM) has emerged, which enables long-term and undisturbed observation of animals in their living environment, facilitating the capture of a more diverse behavioral repertoire (10–14). Among AHCM approaches, RFID has become a widely adopted technology for reliable individual identification and precise behavioral tracking in group-housed environments (15).

RFID is a non-contact communication technology widely used for the identification and tracking of physical objects and living organisms across diverse fields, including logistics and livestock management (16–19). A typical RFID system consists of a small RF tag containing a memory and unique ID, a reader/writer equipped with antennas, and a host computer. The reader/writer transmits an electromagnetic signal via its antennas, and the tag responds by transmitting data back, enabling reciprocal contactless communication.

For identification of animals, compact, biocompatible glass-encapsulated passive tags are commonly used. They operate without internal batteries by harvesting power from the reader’s electromagnetic field. These highly durable tags are implanted with minimal invasiveness under anesthesia (20–22).

When multiple RFID antennas are deployed within an animal living environment, the system not only identifies each animal’s ID but also determines its spatial position based on which antenna detects the signal. This approach had been employed in pioneering field studies using wild mice since at least the 1990s (23).

Antennas used for typical animal tracking can be broadly classified into two types. The first type, exemplified by the TraffiCage (developed by NewBehavior AG, Zurich, Switzerland), employs an array of tile-type antennas placed typically beneath the housing cage floor to track animal positions in the cage (Figure 1A) (24–28). The second type, exemplified by those used in IntelliCage (developed by NewBehavior), employs ring-type antennas typically fitted around tubes, such as those connecting cages or serving as entrances to specific areas, enabling individual identification as animals pass through (Figure 1B) (29–33).

**Figure 1.**
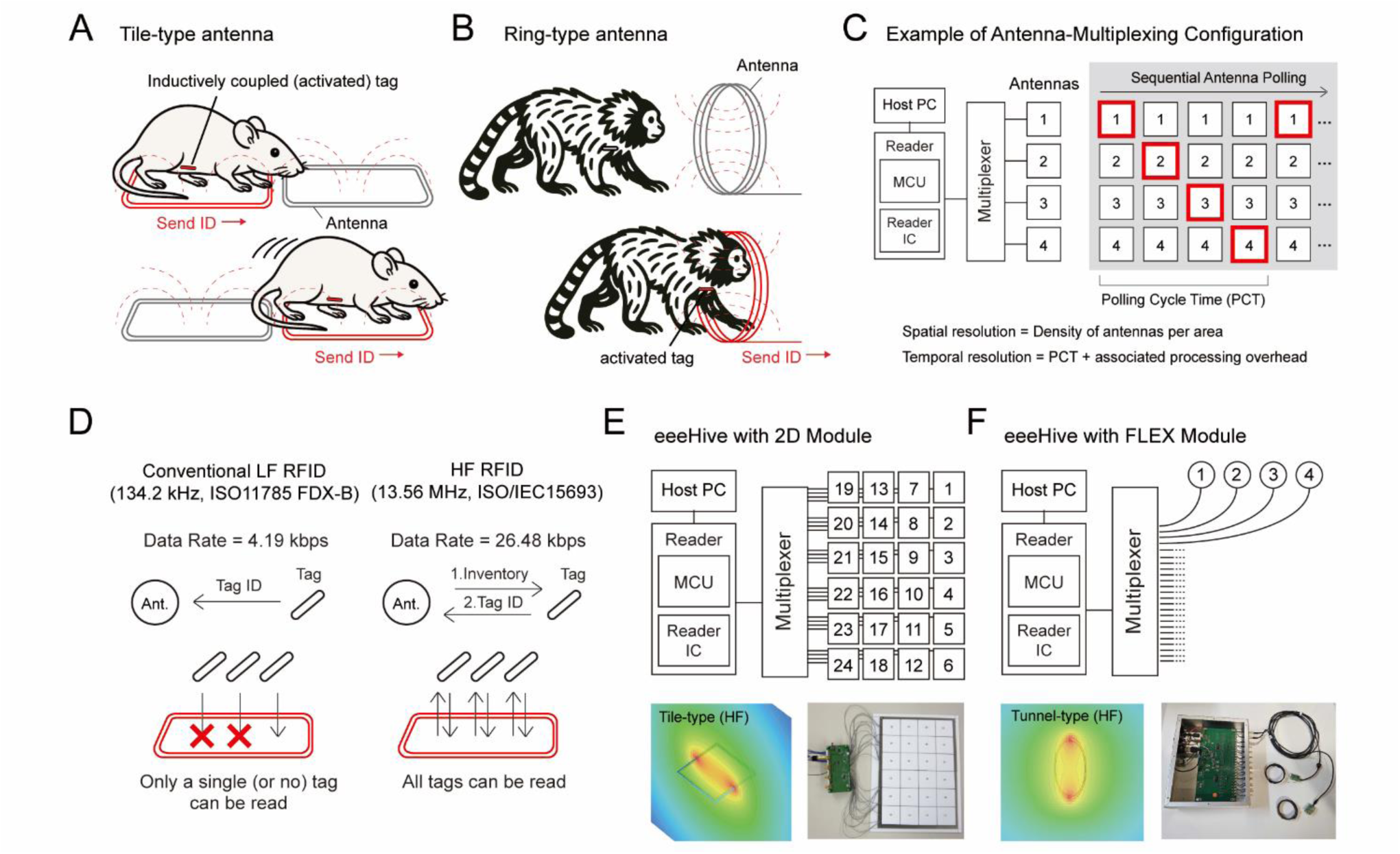
Basics of RFID-based animal behavioral tracking, comparison of LF and HF RFID, and the system architecture of eeeHive with 2D and FLEX modules. (A, B) RFID-based animal behavioral tracking using floor-mounted tile-type antennas (A) and ring-type antennas (B). When the implanted passive tag enters the antenna’s activation field, where the alternating magnetic field (red dotted line) is present, inductive coupling provides the power to transmit the tag’s ID, enabling continuous tracking of animal positions and movements. (C) Schematic diagram of the antenna-multiplexing configuration, composed of host pc, reader, multiplexer, and antennas. To avoid mutual interference, the reader rapidly and sequentially activates each antenna (indicated in red) to detect tags, which is referred to as sequential antenna polling. The time required for a complete scan is defined as the polling cycle time (PCT). The system’s temporal resolution for animal tracking depends on the PCT and the associated processing overhead, while the spatial resolution is determined by the antenna density. (D) Comparison of the conventional LF RFID standard and the HF RFID standard (ISO/IEC 15693). They differ primarily in data transfer rates, communication protocols, and tolerance for tag collisions. LF RFID systems operate at low data rates and use a Tag Talk First (TTF) protocol, causing activated tags to continuously broadcast their IDs and preventing the simultaneous detection of multiple tags by a single antenna. Conversely, HF RFID achieves higher data rates and utilizes a Reader Talk First (RTF) protocol, in which tag responses are strictly controlled by reader commands. This allows the HF system to effectively implement anti-collision mechanisms, enabling the simultaneous reading of multiple tags within the same antenna field. (E) The eeeHive 2D module consists of a modular board containing 24 tile-type antennas (50 × 50 mm each) arranged in a 4 × 6 grid and connected to a single reader. Multiple boards can be seamlessly tiled to expand the tracking area and centrally connected to a single host computer via a USB hub. The lower panel shows the simulated magnetic field strength distribution around a tile antenna and a photograph of the hardware components. (F) The eeeHive FLEX module allows up to 24 custom-shaped and sized antennas per reader with varying cable lengths. These antennas can be freely positioned in three-dimensional space. In this study, ring-type antennas were used for detecting passage events and zone transitions. The lower panel shows the simulated magnetic field distribution of a ring-type antenna and a photograph of the hardware components.

From an ethological standpoint, the multi-antenna RFID tracking approach offers four key advantages. First, it ensures reliable identification of individuals and fine localization of their positions in group-housed environments (31,33). Second, unlike optical systems, it operates independently of lighting conditions and reliably detects visually obscured tags through various media, including sand and soil (34–36). Third, it facilitates large-scale and long-term experiments, as multiple antennas can be cost-effectively deployed throughout the environment while imposing minimal computational and data storage loads over time (30,31,37). Fourth, this approach generates structured behavioral data that are inherently compatible with database systems, thereby enabling efficient querying, analysis, and visualization (15).

In multi-antenna RFID systems, simultaneous activation of adjacent antennas can cause electromagnetic field overlap and mutual coupling, disrupting reader–tag communication and leading to signal collisions and corrupted tag responses. To avoid this issue, such systems typically adopt one of three configuration types described in Supplementary Table 1. Among them, Figure 1C illustrates an antenna-multiplexing configuration in which RF fields are sequentially activated for individual antennas to enable detection of nearby tags. The time required to complete one full round of polling across all connected antennas is defined as the polling cycle time (PCT), and the PCT divided by the total number of antennas corresponds to the polling time per antenna.

High spatial and temporal resolutions are required to reliably detect animal movements within an enclosure. In configurations employing sequential polling, spatial resolution is determined by the antenna density within the enclosure, whereas temporal resolution depends on the PCT and the associated processing overhead. These temporal constraints are largely governed by the data rate, which depends on the RFID frequency.

Importantly, however, the low-frequency (LF) RFID employed in current multi-antenna animal tracking systems (typically operating at 134.2 kHz under ISO 11784/11785 FDX-B protocols) presents several technical limitations (24,28,32,38). Specifically, LF RFID-based system is constrained by its inherent low data rates, which lead to two critical limitations: (i) compromised spatiotemporal resolution, and (ii) the inability to identify multiple tags simultaneously (39–41) (Figure 1D, left). These limitations significantly hinder the analysis of animals like mice that move rapidly and aggregate closely.

Regarding the first limitation in LF RFID, the ISO 11784/11785 FDX-B, for example, is limited by its 4.19 kbps data rate, which requires at least 30.52 ms to receive a full data block from a tag. Additionally, when switching between multiple antennas, the reader must continuously generate an activation field for this entire duration, plus additional processing time for each antenna. Critically, because the reader cannot verify the absence of a tag, this full activation time is completely wasted on empty fields.

In contrast, high-frequency (HF) RFID, typically operating at 13.56 MHz, has a much higher data rate (Figure 1D, right) (42–44). For example, in ISO/IEC 15693, the data rate is 26.48 kbps, requiring only ∼5.9 ms to request and receive a tag ID. Furthermore, if no tag is present, the reader identifies the empty field in ∼2.1 ms and immediately advances to the next antenna. Crucially, this rapid polling enables denser antenna placement within the monitoring enclosure, thereby improving both temporal and spatial resolution.

Regarding the second limitation, in conventional LF RFID, the simultaneous activation of multiple tags within a single antenna field results in mutual signal interference, a phenomenon known as tag collision, which causes critical data loss. This inability to resolve concurrent transmissions severely compromises precise animal localization, particularly in species that naturally engage in dense aggregations and coordinated movements.

Conversely, HF RFID protocols utilize anti-collision functions that divide the response period into discrete time slots (44,45). This temporal separation prevents signal overlaps, allowing the reader to accurately and simultaneously identify multiple individuals huddled together (Figure 1D, right).

Despite these advantages of HF RFID over conventional LF RFID, to the best of our knowledge, no AHCM system leveraging this technology has yet been implemented in practice for a variety of laboratory animals. Therefore, in this study, we sought to evaluate the scientific validity and feasibility of HF RFID by developing a novel animal behavioral tracking system compatible with ISO/IEC 15693 standard (42,44,46). We named our system eeeHive, an acronym for ’easy extensions for ethologically-relevant and home cage-integrated *in vivo* experiments’.

Using this eeeHive, we conducted experiments in C57BL/6J mice (*Mus musculus*), common marmosets (*Callithrix jacchus*; hereafter marmosets), Japanese fire-bellied newts (*Cynops pyrrhogaster*; newts), and juvenile Nile tilapia (*Oreochromis niloticus*; tilapia).

## 2 Materials and Methods

### 2.1 System Overview: eeeHive

#### 2.1.1 Tools for System Development

The eeeHive is a system with a fully custom hardware architecture, designed, assembled, and tested by Phenovance LLC (Chiba, Japan) in collaboration with several subcontractors, as detailed below. The electronic circuit boards and RFID antennas were designed using Altium Designer (Altium Limited, California, USA) and their impedance characteristics were optimized using a network analyzer. Firmware development was performed using Code Composer Studio IDE (Texas Instruments Inc., Texas, USA). The printed circuit boards were fabricated by Showa Sangyo Co., Ltd. (Tokyo, Japan). The physical enclosure was designed with the aid of SOLIDWORKS (Dassault Systèmes SolidWorks Corporation, Massachusetts, USA). A model of the magnetic field strength distribution was constructed by Dr. Hiroaki Kogure (Kogure Consulting Engineers, Tokyo, Japan) using S-NAP® Wireless Suite software (MEL Inc., Nagoya, Japan), based on the CAD data of the reader antenna.

#### 2.1.2 HF RFID Principle, Protocols, and Tags

The eeeHive system employs HF RFID. Several international standards (ISO) for HF RFID communication include ISO/IEC 14443 Type A and B, ISO/IEC 15693, and ISO/IEC 18000-3. Based on preliminary evaluation, we adopted ISO/IEC 15693 for the eeeHive, which provides a longer read range than other HF standards. Table 1 summarizes a comparison of the key parameters between ISO 11784/11785 FDX-B and ISO/IEC 15693.

**Table 1.** Comparison of Key Specifications of LF RFID (ISO 11784/11785 FDX-B) and HF RFID (ISO/IEC 15693)

| Key specifications | LF RFID compliant with<br>ISO 11784 / 11785 FDX-B | HF RFID compliant with<br>ISO/IEC 15693 |
| --- | --- | --- |
| Year of publication | ISO 11784: 1st ed. 1994-05, 2nd ed. 1996-08, 3rd ed. 2024-12; ISO 11785: 1st ed. 1996-10 | 1st ed. 2000 (Parts 1-3); current 3rd ed.: Part 1: 2018, Parts 2&3: 2019 |
| Operating frequency | 134.2 kHz | 13.56 MHz |
| Coupling mechanism | Inductive coupling | Inductive coupling |
| Data transfer rate | 4.19 kbps (fc/32) | 26.48 kbps (high-speed); 1.65 kbps (low-speed) |
| Modulation | Backscatter ASK via load modulation | Reader to tag: ASK (10% or 100% depth); Tag to reader: Load modulation with subcarrier (ASK or FSK) |
| Data encoding | Differential biphase (modified) | Reader to tag: Pulse-position coding (1-of-4 or 1-of-256); Tag to reader: Manchester coding |
| Communication sequence | Tag talks first (TTF): Tag responds during reader excitation | Reader talks first (RTF): Tag responses are controlled by Reader |
| Tolerance to tag collisions | Not supported | Supported (slotted ALOHA, 16 fixed slots) |
| Penetration depth - freshwater (0.005–0.15 S/m) | 3200–26,000 cm* | 30–2,500 cm* |
| Penetration depth - seawater (4.0 S/m) | 71 cm* | 6.8 cm* |
| Memory architecture | Fixed 128-bit telegram | Flexible block-structured memory |
| Standardization scope | Dedicated to animal identification | General-purpose contactless vicinity objects |

\*These values are based on Benelli and Pozzebon (2013). Note that the LF figures strictly refer to 125 kHz tag data.

Across all tested species, cylindrical biocompatible glass-encapsulated tags (2.12 mm in diameter, 12 mm in length) with an ISO/IEC 15693-compliant chip (EM Microelectronic, Switzerland), commercially available from Phenovance LLC (Chiba, Japan), were used.

#### 2.1.3 Hardware Overview

##### 2.1.3.1 General Architecture

The eeeHive system adopted the antenna-multiplexing configuration. It consists of a reader with an antenna multiplexer, multiple reader antennas, and a host computer. The reader was designed to generate a stable 13.56 MHz magnetic field, manage antenna multiplexing, and handle communication with passive tags. The reader antennas were impedance-matched and designed to generate a uniform magnetic field throughout the detection area. The reader includes a microcontroller unit (MCU; Texas Instruments, Dallas, TX, USA), which interfaces via SPI with an RFID reader IC (NXP Semiconductors, Eindhoven, Netherlands), and a connector for transmitting control signals to the antenna multiplexer. The antenna multiplexer uses RF switches (pSemi Corporation, San Diego, CA, USA) to sequentially activate up to 24 connected reader antennas based on control signals received from the reader. The reader was connected to the host computer via a UART-to-USB converter. A laptop computer (Lenovo ThinkPad X280) was used as the host system for data acquisition and control.

##### 2.1.3.2 Two Types of System Configuration with “2D module” and “FLEX module”

Two types of hardware configurations, referred to as the “2D module” and “FLEX module”, were developed for the eeeHive system. Each module comprises a reader, a multiplexer, and multiple reader antennas.

The 2D module was developed for two-dimensional behavioral tracking of animals with high spatial resolution. It consists of a 200 × 300 mm board containing 24 tile-type coil antennas (each 50 mm square) arranged in a 4×6 grid (Figure 1E). Each antenna is assigned an antenna ID (1–24) and connected via coaxial cables to a multiplexer. When placed beneath a cage, the system records animal positions by linking timestamp (millisecond resolution), antenna ID, and tag ID. Multiple boards can be arranged side by side without gaps to expand the tracking area.

The FLEX module permits flexible antenna placement, thereby enabling broader spatial coverage and three-dimensional behavioral tracking (Figure 1F). Up to 24 antennas can be connected to a reader via coaxial cables. In this study, ring-type antennas with diameters of 100 mm for marmosets and 50 mm for tilapia were used. Animal identification occurred when an animal passed through a tunnel. These antennas were placed at key locations such as passages, feeders, water stations, and nestbox entrances to detect behavioral events and spatial transitions.

##### 2.1.3.3 Evaluation of Tag Read Range and Processing Speeds

The tag read range and processing speeds were evaluated for the eeeHive 2D and FLEX modules after optimizing the antenna output parameters (Figure 2).

**Figure 2.**
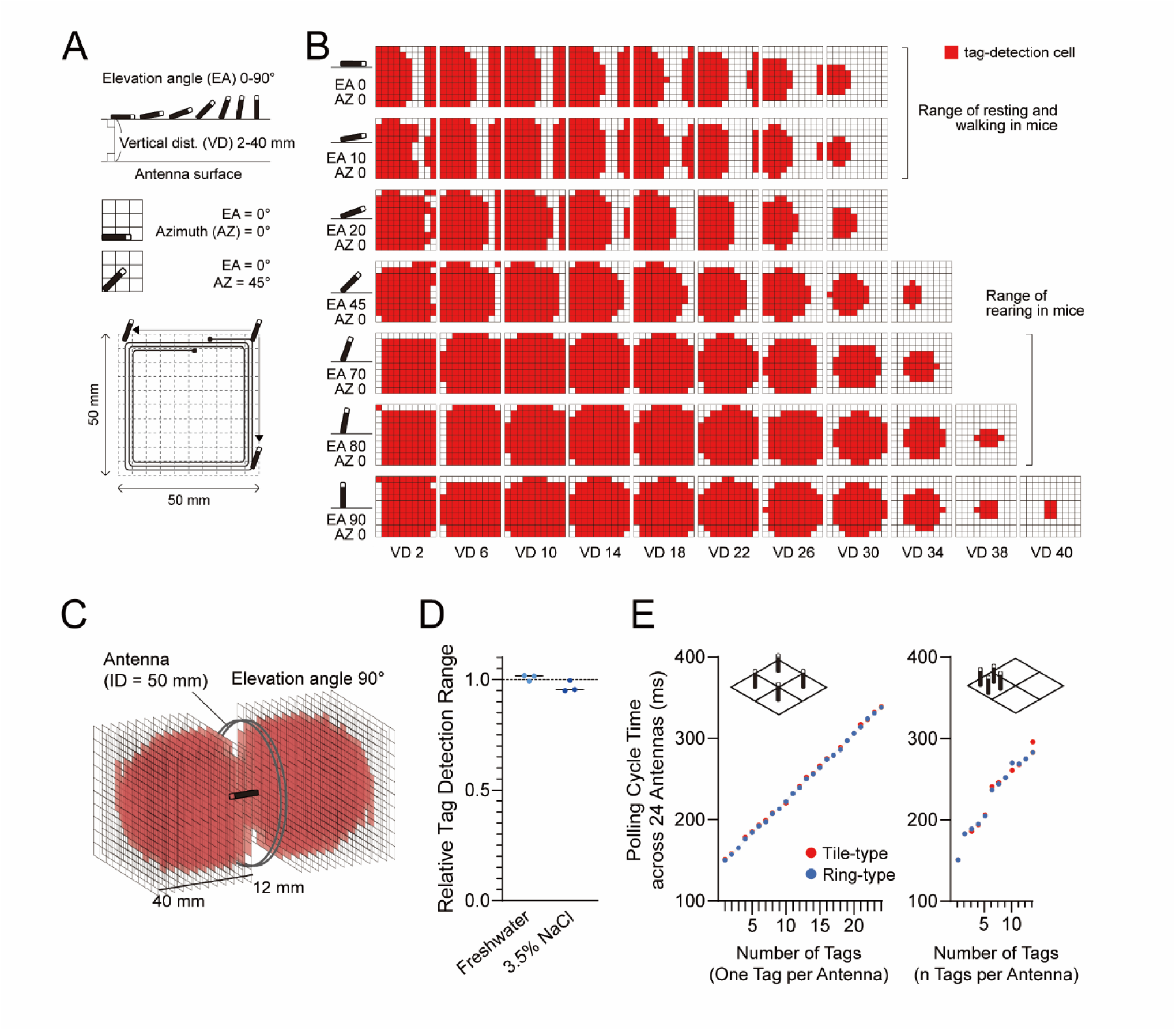
Tag detection range and reading speed of the eeeHive 2D and FLEX modules. (A) Evaluation of tag readability for the tile-type antenna (eeeHive 2D). Readability was assessed by systematically varying the tag’s elevation angle (EA; 0 to 90°), azimuth angle (AZ; 0° and 45°), vertical distance (VD) from the antenna surface (2 to 40 mm), and lateral position across a 50 × 50 mm grid. A total of 15,400 combinations were tested, and successful tag ID detection was recorded for each condition. (B) Spatial distribution of successful tag reads at AZ 0°. Cells with successful detections are indicated in red. Overall, the detectable coverage area decreases with increasing VD. EA between 0° and 10° correspond to the typical range observed when a mouse is resting or walking, whereas angles between 70° and 80° correspond to rearing behavior. Larger EA yields longer readable distances, reaching a maximum of 40 mm. Although the read sensitivity of these antennas is adjustable, it was optimized for conditions simulating mouse tracking. (C) Tag readability of the ring-type antenna (eeeHive FLEX) measured assuming a tag passing perpendicularly through the antenna (EA 90°). Readability was assessed on both lateral sides of the antenna. As in the tile-type configuration, the detectable region exhibited a dome-shaped profile, with a maximum readable distance of approximately 40 mm under the experimental output and sensitivity settings. (D) Maximum tag reading distances in freshwater and a seawater-equivalent 3.5% NaCl solution. Results from three independent antennas are plotted as normalized read ranges relative to the performance in air (Air = 1). (E) Polling cycle time (PCT) and anti-collision cost in the 2D (red) and FLEX (blue) configurations. Left: Total PCT as a function of the number of antennas each containing one tag (up to 24 antennas). PCT increased linearly with the number of tags, with an additional cost of 8.2 ms per tag. Right: Effect of multiple tags within a single antenna field. Nonlinear increases in PCT were observed with increasing tag count, reflecting the computational cost of anti-collision processing (approximately 30 ms for the first few tags).

Tag readability was assessed across 15,400 spatial conditions near a tile-type antenna, combining seven elevation angles, two azimuth angles, 100 horizontal positions (Figure 2A), and 11 vertical offsets. Detection results for the 0° and 45° azimuths are presented in Figure 2B and Supplementary Figure 1A, respectively. The same procedure was applied to the ring-type antenna, except that cells corresponding to regions outside the circular coil were excluded from the analysis.

All evaluations were performed experimentally according to the procedures described in the Supplementary Information 1.

### 2.2 System Implementation per Species

Animal experiments were conducted at Phenovance LLC (mice, newts, tilapia) and Hamamatsu University School of Medicine (marmosets), with the approval of the relevant Institutional Animal Care and Use Committees (Experiment Approval Numbers: PRT202402_48_2 for Phenovance LLC and 2020058 for Hamamatsu University) and in accordance with the applicable regulations, guidelines, and 3R principles regarding the conduct of animal experiments.

#### 2.2.1 Mice

We used a cohort of 24 male C57BL/6J mice (M01–M24), aged 17 months and weighing over 29 g, that had previously served as controls in other experiments. They were offspring of dams obtained from The Jackson Laboratory Japan (Yokohama, Japan) and had participated in multiple behavioral experiments prior to the present study, without prior exposure to invasive procedures or pharmacological interventions.

A tag was implanted subcutaneously along the ventral midline of the abdominal area using a syringe under general anesthesia induced by 3% isoflurane (Myland Pharmaceuticals Co., Ltd., Japan) delivered via a precision vaporizer (SN-487-1T Inhalation System, Shinano Seisakusho Co., Tokyo, Japan).

Based on previous observations indicating a low incidence of inter-male aggression at this age, mice were group-housed in a custom-designed acrylic cage (40 × 60 × 13 cm) constructed from 5 mm-thick transparent, colorless acrylic panels bonded using an acrylic-specific adhesive. The floor area and height of the enclosure were within internationally accepted guidelines for group-housed mice exceeding 25 g in body weight (47,48).

Four eeeHive 2D modules were in a 2 × 2 grid arrangement, beneath the cage (Figures 3A and 3B). They were connected to a host PC via a single USB hub. The cage was subdivided into five compartments (Rooms 1–4 and Center) by dividing walls, allowing free movement between compartments. The cage lid consisted of two panels (40 × 30 cm each), one transparent and colorless and the other transparent red, resulting in a half-clear and half-shaded enclosure. The sidewalls, dividing walls, and lids were mechanically perforated with arrays of 10 mm-diameter apertures spaced at 3–5 cm intervals.

**Figure 3.**
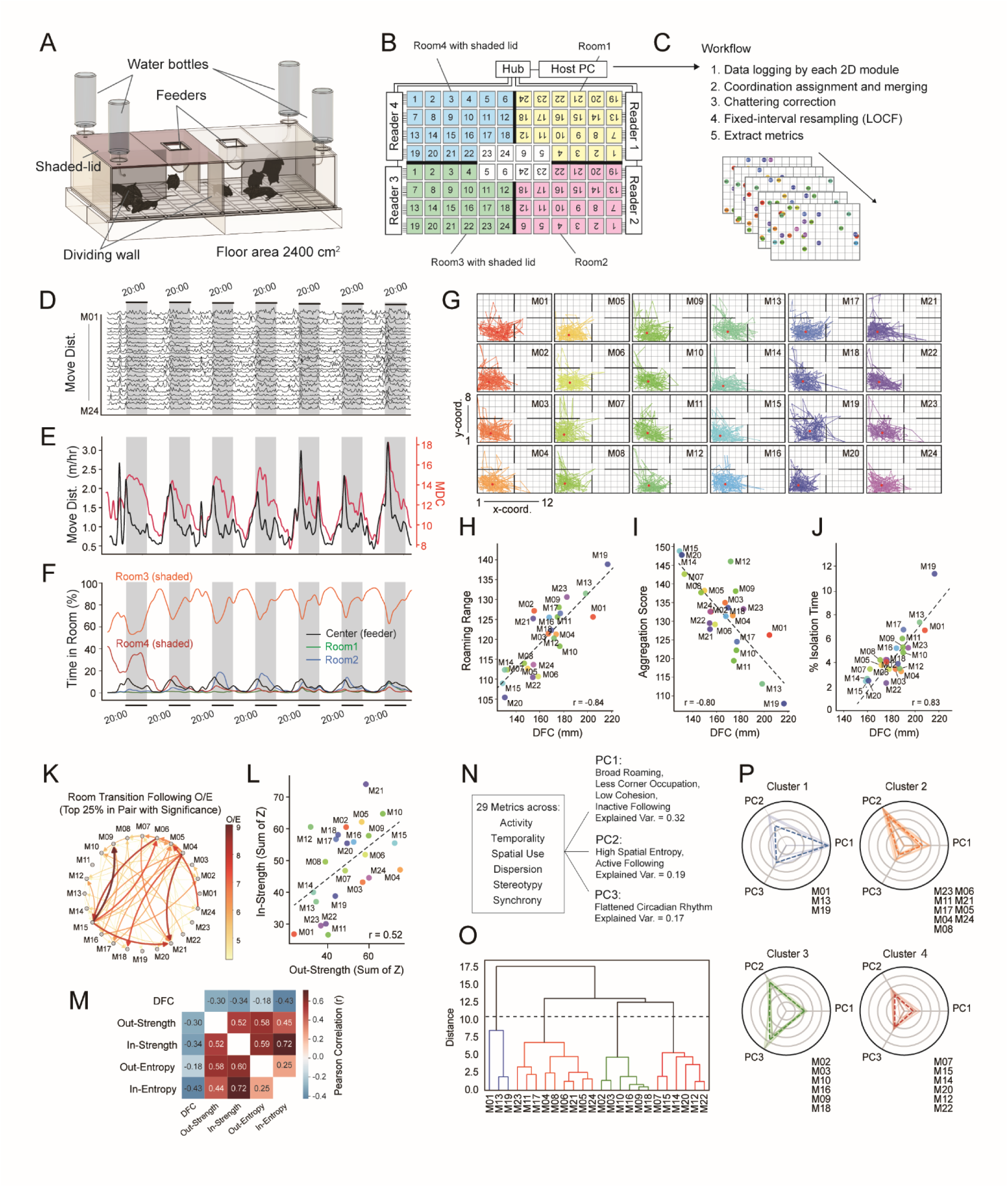
Behavioral analysis of group-housed mice using the eeeHive 2D module. (A) Illustration of the experimental enclosure. Twenty-four mice were group-housed in a custom-designed acrylic cage (40 × 60 × 13 cm) placed on four eeeHive 2D modules. The cage floor area (2,400 cm²) fully covered the 8 × 12 antenna grid. Dividing walls separated the interior into five compartments (Rooms 1–4 and the Center), and mice were able to freely transition between compartments through the central area. The ceilings of Rooms 1 and 2 were made of transparent acrylic plates, whereas those of Rooms 3 and 4 were made of red acrylic plates, creating differences in illumination between the two areas. (B) Configuration of the setup. Four eeeHive 2D modules were connected to a host computer via a self-powered USB hub. This configuration enabled the simultaneous acquisition of tags from 96 antennas while maintaining the polling cycle time (PCT) of a single 2D module. Rooms 1 to 4 are shown in yellow, pink, green, and blue, respectively, and the Center in white. (C) Workflow of data preprocessing and analysis. (D) Activity profiles of all 24 mice across seven days. The x-axis represents time, with dark phases shaded in gray. The y-axis indicates the distance traveled. Individual traces are vertically stacked for visualization. Activity was quantified as the cumulative movement distance per hour. Movement distances were calculated based on changes in antenna coordinates (Δx, Δy), assuming a center-to-center distance of 50 mm between adjacent antennas and 50 √ 2 mm for diagonal displacements. (E) The black trace indicates mean movement distance across all individuals (m/h), and the red trace indicates the Distance from Centroid (MDC), the mean distance from each individual to the group’s spatial centroid (cm) at each time point, representing group dispersion. (F) Temporal changes in the percentage of time spent in each compartment. Values represent the mean proportion of time each individual spent in each compartment during each one-hour period. The shaded background indicates the dark phase. (G) Hourly centroid trajectory plot for each individual across the 7-day experimental period. The red dot indicates the individual global centroid (IGC), defined as the individual’s spatial centroid across the entire recording duration. (H) Relationship between roaming range and distance from the corner (DFC). Roaming range was defined as the mean distance (mm) between each individual’s sampled coordinates and its IGC across the entire experimental period. DFC was defined as the mean distance (mm) between each individual’s sampled coordinates and the Room 3 corner reference point (coordinate 1,1). r = 0.84, p < 0.05. (I) Relationship between DFC and Aggregation Score, defined as the cumulative co-presence time on the same antenna (50 mm × 50 mm), weighted by the number of co-present individuals and the duration of overlap. r = −0.80, p < 0.05. (J) Relationship between DFC and isolation time (%), defined as the proportion of the total experimental duration during which no other individual was present on the same antenna or on any of the four immediately adjacent antennas. r = 0.83, p < 0.05. (K) Directed network of the follow behavior with observed over expected (O/E) ratio as edges. Edges represent the top 25% of O/E ratios. (L) Relationship between out-strength as the sum of Z-scores for follow behaviors directed from a given individual to all other individuals, and in-strength as the sum of Z-scores for follow behaviors received from all other individuals. r = 0.52. (M) A correlation matrix across five metrics: DFC, out-strength, in-strength, out-entropy, and in-entropy. (N) PCA of 29 behavioral metrics encompassing activity, temporality, spatial use, dispersion, and stereotypy. (O) Hierarchical clustering of individuals based on the 29 behavioral metrics. The dendrogram (Ward’ s method) reveals four distinct clusters, indicated by color coding. The dashed horizontal line represents the clustering threshold used to define the four groups. (P) Radar plots illustrating the relative contribution of PC1–PC3 scores for individuals classified into four behavioral clusters.

Two feeders (8 × 5 × 6.5 cm each) were mounted on the ceiling, positioned such that feeding occurred entirely within the center area. Standard rodent chow (Labo MR Standard, NOSAN Nosan Co., Ltd., Kanagawa, Japan) was provided. Water bottles (250 mL-size each) were installed at the four corners of the cage. Neither food nor water required replenishment during the 7-day data-acquisition period. The animal facility was maintained under a 12-hour light–dark cycle (lights off: 20:00–08:00), with a room temperature of 25 ± 2°C and a relative humidity of 40–70%.

#### 2.2.2 Marmosets

The marmoset colony comprised 11 marmosets of both sexes and various ages. Tags were implanted subcutaneously in the dorsal neck region of six animals (four males M1–4; two females F1–2), and only these tagged individuals were included in the behavioral analysis. Two of the analyzed males (M1, M4) were bred in-house, whereas the others were obtained from CLEA Japan (Tokyo, Japan). For tag implantation, animals were anesthetized with a combination of medetomidine (Domitor, Nihon Zenyaku Kogyo Co., Ltd., Fukushima, Japan), midazolam (Dormicum, Astellas Pharma Inc., Tokyo, Japan), and butorphanol (Betulfa, Meiji Animal Health Co., Ltd., Kumamoto, Japan). The insertion site was closed with a single nylon suture.

Marmosets were housed in an enclosure with a total internal volume of 44.65 m³ (4.7 × 3.8 × 2.5 m). One wall was fitted with a 1 cm × 1 cm wire mesh grid, which served as the primary access point for caretakers and experimenters. A lockable door within the mesh provided controlled entry, and a vestibule outside the mesh created a double-door system for enhanced containment and environmental control. The enclosure was enriched with a jungle gym comprising 10 wooden poles (2–3 m in length, 100 mm in diameter), wooden perches, ropes, bridges, nets, hammocks, and nestboxes.

Nestboxes varied in size as follows: one large (Nest 2; 33.5 × 48 × 22 cm) accommodating 8–10 animals; two medium (Nests 1 and 3; 24 × 35 × 18 cm and 25 × 34.5 × 15 cm) for 6–8 animals; one small (Nest 4; 14 × 21 × 14 cm) for 2–3 animals; and two extra-small (Nests 5 and 6; 14 × 16.5 × 18 cm) for 1–2 animals. Nests 5 and 6 were rarely used and were therefore excluded from the analysis.

A total of 20 ring-type FLEX antennas (inner diameter: 100 mm) were distributed throughout the enclosure: one antenna at each of the four water stations and four feeders, and two antennas at each of the six nestboxes (Figure 4B). At each nestbox, a pair of antennas was installed at the entrance tube, 50 mm apart, with the one closer to the nestbox designated as the inner antenna and the other as the outer antenna. When an animal passed through the entrance tube, the direction of its movement was determined based on the time difference in tag detection between the two antennas. A ferrite sheet was inserted between the two antennas to prevent simultaneous tag detection.

**Figure 4.**
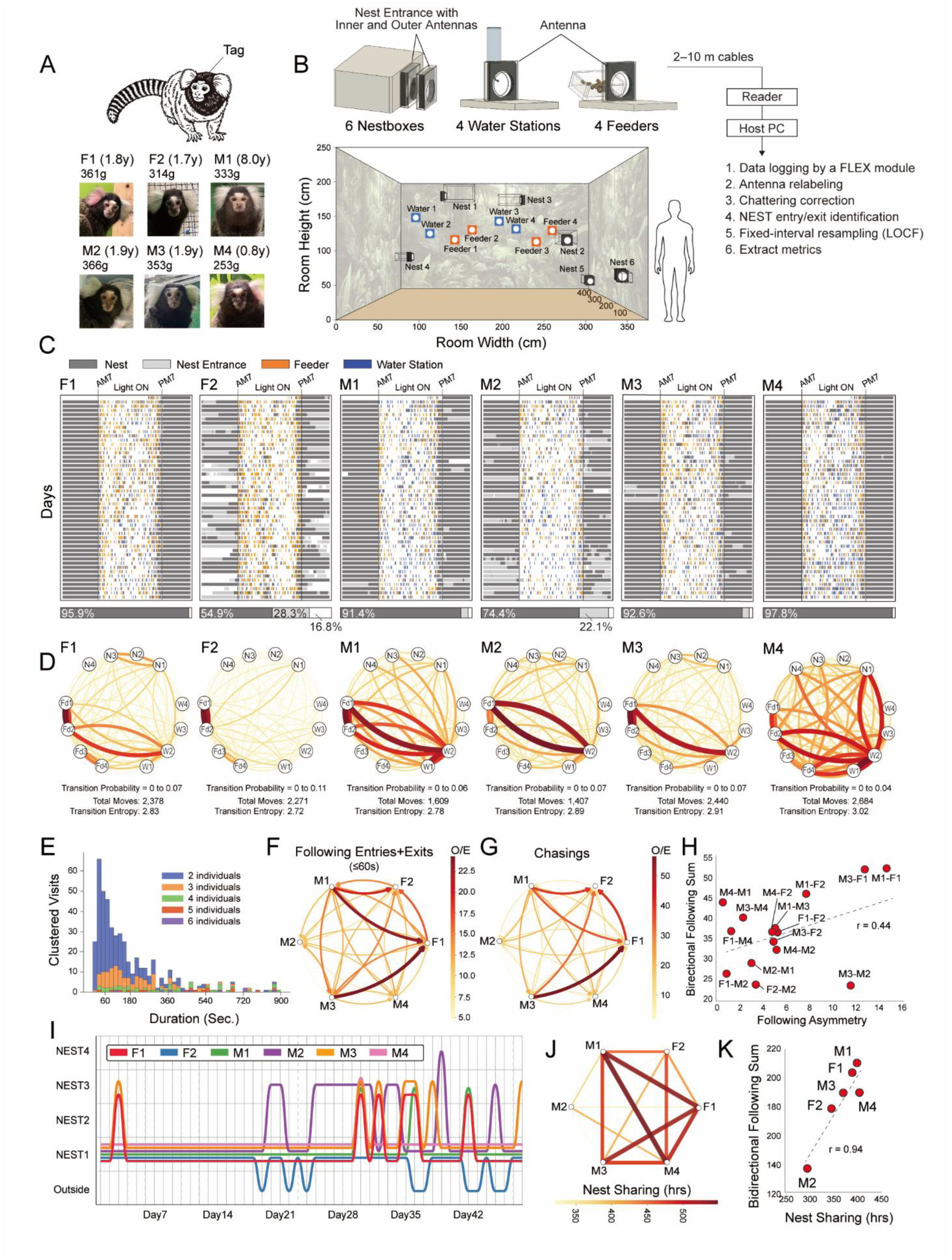
Behavioral analysis of group-housed marmosets using the eeeHive FLEX module. (A) Tags were subcutaneously implanted in the nuchal region of six marmosets. Their respective names (F1, F2, M1, M2, M3, M4), ages, and average weights during the experimental period are shown. (B) Schematic illustration of the experimental setup. Marmosets were housed in an enclosure of approximately 44.65 m³ (4.7 × 3.8 × 2.5 m) with various enrichment structures and accessories (Not shown in the illustration; see Materials and Methods for the detailed description). Six nestboxes (Nest 1–6) of varying sizes, four water stations, and four feeders were each equipped with a FLEX module. For each nestbox, a pair of antennas (outer and inner) was installed at the entry/exit points to enable directional detection of entries, exits, and dwelling behavior. (C) Visualization of location logs of each individual across seven weeks. The x-axis represents time of day (leftmost = 00:00), and the y-axis represents experimental days. The color at each time point indicates the location where the individual was detected. White indicates periods during which the individual was not detected at any monitored location. (D) Visualization of transition probabilities among four main nestboxes (Nest 1–4), two water stations (Water 1–2), and two feeders (Food 1–2) for each individual. Transitions with higher probability are depicted in darker red and with thicker edges. Of the two edges connecting a pair of nodes (locations), the edge curving to the right indicates the direction of movement. For each individual, the minimum and maximum transition probabilities used for the color scale, the total number of transitions, and first-order transition entropy are indicated below the networks. (E) Clustered visits recorded during the light phase. The x-axis represents the duration from the start to the end of each clustered visit, and the y-axis represents frequency. Colors indicate the number of individuals per clustered visit. (F) Directed network based on O/E ratios of the following events (entries and exits) within each pair. (G) Directed network graph based on O/E ratios of bidirectional chasing within each pair. (H) Scatter plot showing the relationship between asymmetry in follow relationships and the total amount of bidirectional follow behavior within each pair. For bidirectional Following Entries + Exits, Z-scores obtained from permutation tests were used; the difference between the two individuals’ Z-score difference is plotted on the x-axis, and their sum on the y-axis. In the pair labels shown in the figure, the individual with the higher Z-score for Following Entries + Exits is listed first. r = 0.44. (I) Visualization of the primary location used by each individual during the dark phase of each experimental day. The x-axis represents experimental days (Day 1–48), and the y-axis represents location. (J) Network graph with nestbox co-occupancy time for each pair represented as edges. (K) Scatter plot showing the relationship between each individual’s mean nocturnal nestbox co-occupancy time with others (x-axis) and the total bidirectional following (sum of bidirectional Following Entries + Exits Z-scores; y-axis). r = 0.94.

The colony was maintained under a 12-hour light–dark cycle (lights off: 19:00–07:00), with temperature regulated at 25 ± 3 °C and relative humidity at 40–70%. Animals were maintained under a standardized feeding regimen, with details provided in Supplementary Information 2. During daily husbandry routines (∼2 h each morning), three experienced animal caretakers monitored health status and recorded behavioral observations (Supplementary Information 3), with individuals identified based on physical features and color markings.

#### 2.2.3 Newts and Tilapia

Four newts (2 males M1–2, and 2 females F1–2; approximately 100 mm in length) and eleven juvenile tilapia (M1–11; 50–70 mm in length) were obtained from a local supplier in Japan. Intracoelomic implantation was conducted using a syringe, under anesthesia with 0.1% ethyl 3-aminobenzoate methanesulfonate (MS-222; CAS: 886-86-2; Sigma-Aldrich Co., LLC, USA). The wound was closed using a medical tissue adhesive (derma+flex, Chemence Medical, Inc., Georgia, USA).

For housing, custom-made, species-specific acrylic tanks were constructed from transparent acrylic panels. For the newt study, a tank (24 L; 60 × 40 × 15 cm; 3 mm base and 5 mm wall thickness) filled with 1 cm-deep freshwater was placed on two eeeHive 2D modules (8 × 6 array; 48 antennas total) (Figure 5A). For the tilapia study, a tank (25 L; 60 × 20 × 25 cm; 5 mm wall thickness) was divided into two compartments connected by a tunnel (inner diameter ∼45 mm). A pair of ring-type antennas was installed, one at each end of the tunnel, separated by a 50 mm gap (Figure 5G). A ferrite sheet was inserted between them to prevent simultaneous tag detection. The antennas were waterproofed and were fully submerged in freshwater.

**Figure 5.**
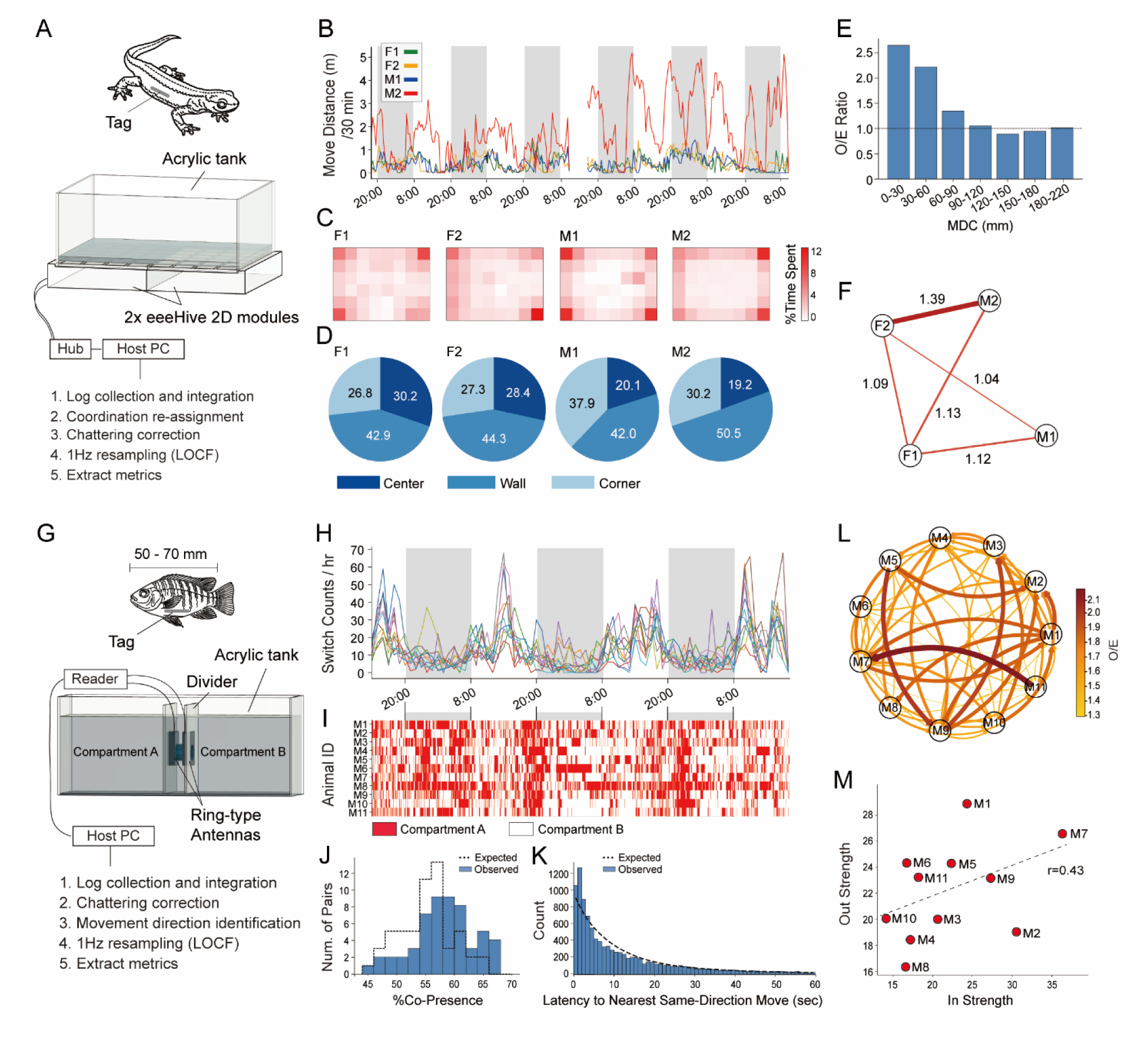
Application of the eeeHive system to behavioral analysis in aquatic species. (A) Experimental setup and data-processing workflow for the newts experiment. A water-filled acrylic tank (60 × 40 × 15 cm) was placed on top of two 2D modules. The tank was filled with water to a depth of 10 mm. (B) Time series of movement distance per 30-min bins across the experimental period. The y-axis indicates the distance travelled per 30 min (m), and the x-axis indicates time. Dark phases (20:00–8:00) are shaded. (C) Heatmaps showing the proportion of time each newt spent at each location within the tank across the entire experimental period. A common color scale (with a shared maximum) was applied to all individuals to enable direct comparison. (D) Proportion of time spent in each spatial zone category (corner, wall, and center) for each newt, expressed as a percentage of total stay time. (E) The mean distance from centroid (MDC; mm) among the four newts was calculated at 1-s resolution and binned into distance ranges. For each bin, the O/E ratio based on 1,000 permutation null datasets is shown. The MDC between 0 and 90 mm showed significantly elevated O/E ratios (p < 0.05), with O/E ratios increasing progressively in smaller MDC bins. (F) Undirected network representation of pairwise proximity relationships (inter-individual distance ≤ 100 mm) based on O/E ratios, illustrating non-random associations among newts. Only edges with statistically significant proximity (permutation test, p ≤ 0.05) are shown. (G) Experimental setup using the eeeHive FLEX module and data-processing workflow for the tilapia experiment. The tank (60 × 20 × 25 cm) was divided into two compartments by a tunnel (inner diameter: 45 mm), with two ring-type antennas installed at each end. The antennas were fully submerged in water and positioned ∼5 cm apart, with a ferrite shield between them to prevent simultaneous tag detection. (H) Hourly trafficking events of tilapia (compartment-switching counts per hour) throughout the experimental period. Dark phases (20:00–8:00) are shaded. (I) Temporal structure of compartment use for each tilapia. Individual locations were reconstructed at 1-s resolution. Compartment A is shown in red and Compartment B in white. (J) Distribution of pairwise co-presence across all 55 tilapia pairs. The observed data are compared with the mean null expectation derived from 1000 permutation trials. The x and y axes represent co-presence (%) and the number of pairs, respectively. (K) Distribution of inter-individual follow latencies (1 s bins). Observed values represent the shortest latency for another individual to move in the same direction after an initial movement. The null distribution was generated from 1,000 permutation trials. The x- and y-axes represent latency (s) and event frequency, respectively. (L) Directed follow network illustrating the directionality of follow interactions, based on O/E ratios. Only significant directed edges are shown (permutation test, p < 0.05). For each pair of individuals, the right-curving edge indicates the direction of movement. (M) Scatter plot of out-strength versus in-strength for each individual in the tilapia group. Out-strength and in-strength were calculated as the sums of Z scores for outgoing and incoming follow edges, respectively. Each point represents one individual. r = 0.43.

The animal facility was maintained under a 12-hour light/dark cycle (lights off: 20:00–08:00) at a room temperature of 23 ± 3 °C. Mechanical filtration systems were installed to maintain water quality. Newts were fed 5 g per day of vitamin-enriched frozen bloodworms (Kyorin Co., Ltd., Tokyo, Japan), thawed in running water and offered every 2–3 days, consistent with the dietary requirements of carnivorous amphibians. Tilapia were fed TetraFin® flake food (Tetra GmbH, Melle, Germany) three times daily.

### 2.3 Data Analysis

#### 2.3.1 Tools

Communication between the eeeHive 2D/FLEX modules and the host computer was conducted using free terminal software Tera Term installed on the host computer. The acquired data were saved as plain text log files, in which each line represented a single detection record consisting of a timestamp, the antenna ID, and a list of one or more tag IDs detected by that antenna. The complete dataset consisted of a sequence of such records. Data preprocessing, analysis, statistics, and visualization were performed using Python (3.11.9) and the following additional libraries: pandas (2.2.2), numPy (1.26.4), matplotlib (3.9.0), seaborn (0.13.2), networkX (3.3), statsmodels (0.14.5), scipy (1.15.1), and scikit-learn (1.7.2). For statistics, graphing, and illustrations, Adobe Illustrator (Adobe Inc., San Jose, CA, USA), and Autodesk Fusion (Autodesk Inc., San Francisco, CA, USA) were used. Animal illustrations were created using licensed Adobe materials.

#### 2.3.2 Data Processing Workflow

##### 2.3.2.1 Workflow for 2D module Experiments

The workflow from data acquisition to analysis in the mouse and newt experiments using the 2D module was as follows (Figures 3C and 5A).

First, log files independently collected by multiple 2D modules were saved to the host PC. The antenna IDs (1–24) in each log file were converted into coordinate data based on the spatial configuration of the corresponding 2D setup. For example, in the mouse experiment, antenna 19 of reader 3 (Room 3) was assigned coordinates (1, 1), whereas antenna 19 of reader 1 (Room 1) was assigned coordinates (12, 8) (Fig. 3B). The separately recorded log files were then merged into a single text file.

Next, data noise processing was performed. When multiple antennas are positioned in close proximity, their tag detection ranges may partially overlap. Consequently, a tag entering the overlapping region may be detected by multiple antennas. Under these conditions, even if the tag remained stationary, sequential polling could generate log entries indicating as if the tag is rapidly moving back and forth between adjacent antennas several times per second. This artifact is referred to here as “chattering”.

To eliminate the chattering artifacts, the following rule was applied. For a given tag detected at antenna i at time t₁, subsequently at one or more other antennas, and again at antenna i at time t₃, if (t₃ − t₁) < Δt, all detections at antennas other than i occurring between t₁ and t₃ were removed from the dataset. Unless otherwise specified, Δt was set to 1 s. For the newt experiment, Δt was set to 2 s based on the distribution of antenna-switching intervals calculated from linear bouts without immediate reversals, which indicated that switching events within 2 s likely reflected detection fluctuations rather than genuine movement.

At this stage, because each line in the log file represented a timestamped detection record, the time intervals between successive records were not uniform and varied on the millisecond scale. To standardize the temporal resolution for subsequent analyses, a new dataset was generated in which the coordinates of each tag were recorded at fixed 1-s intervals (or 0.2 s for certain analyses). This step is hereafter referred to as fixed-interval resampling. The coordinate of each tag at each time point was completed using the last observation carried forward (LOCF) method. From the pre-processed and resampled data, various metrics were calculated, including individual travel distance, activity rhythms, spatial utilization, inter-individual distances, and reciprocal following behavior.

##### 2.3.2.2 Workflow for FLEX module Experiments

The analysis workflow for the common marmoset and tilapia experiments using the FLEX module was as follows (Figures 4B and 5G). Log files collected by the FLEX module were saved to the host PC. The antenna IDs (1–24) in the log files were relabelled according to the physical locations where the antennas were installed.

As in the 2D module, some antennas were installed in close proximity in the FLEX module setup. Therefore, chattering removal was performed using the same method described above. From the processed data, the timing of entry and exit events at the nestbox (marmoset experiment) and inter-compartment transitions (tilapia experiment) was identified.

To standardize the time intervals in the dataset, fixed-interval resampling with LOCF was subsequently performed. Using the resampled data, various behavioral metrics were calculated, including individual movement, activity rhythms, spatial utilization, and reciprocal following behavior.

#### 2.3.3 Permutation Test

To evaluate whether inter-individual behavioral metrics exceeded chance levels, we employed permutation tests. Observed metrics were compared against a null distribution representing entirely independent movements. We generated 1,000 null datasets, primarily using a 10-min block circular shift. This method shifts each individual’s time series by a random offset within 10-min blocks, effectively disrupting inter-individual timing while preserving intrinsic activity rhythms and spatial preferences. By evaluating observed values (O) against the null expected mean (E), we calculated the observed-to-expected (O/E) ratio and Z-score ((O − E) / SD_null) for effect sizes, and p-values for statistical significance.

## 3 Results

Here, we present the performance of tag readability for the 2D and FLEX modules (Figure 2), followed by demonstrations of behavioral tracking in mice (Figure 3) and marmosets (Figure 4), and finally their applicability in aquatic environments (Figure 5).

### 3.1 Performance of Tag Read in eeeHive 2D and FLEX Modules

The tag readable range depends on its spatial orientation and distance from the antenna, while accurately tracking rapid movements requires sufficient polling speeds. Understanding these spatiotemporal constraints of magnetic coupling enables researchers to optimize tag implantation and antenna layout, minimizing missed behavioral recordings.

#### 3.1.1 Spatial Range of Tag Read

We evaluated the tag read range of the eeeHive 2D and FLEX modules (Figures 2A−C, Supplementary Figure 1A) as described in the Supplementary Information 1.

Despite expectations that an elevation angle near 0° (where the tag’s coil opening is orthogonal to the antenna) would hinder detection, coverage remained robust at 76% at a 2-mm vertical distance. As the elevation angle approached a vertical orientation, the readable vertical range increased, and tag IDs could be read at distances of up to 40 mm.

This profile is ideal for abdominally implanted tags in mice. Short distances compensate for parallel orientations during resting or walking, whereas increased elevation angles extend the read range during rearing, effectively minimizing missed detections across varying postures (Supplementary Figure 1B).

We next measured the detection range of the ring-type antenna. To simulate a tag implanted along the body axis passing perpendicularly through the coil, we fixed the elevation angle at 90°. As with the 2D module, the readable range narrowed with distance into a dome shape, reaching a maximum of 40 mm (Figure 2C).

Given that higher-frequency signals are prone to attenuation in conductive media, we tested whether this limits the 40-mm read range. However, the range remained largely unaffected in both freshwater and a 3.5% NaCl solution, exhibiting only a marginal 1.4 mm average reduction in the saltwater (Figure 2D).

#### 3.1.2 Speed of Antenna Polling and Tag Read

We examined the polling cycle time (PCT) of the eeeHive system across 24 antennas as the number of tags increased, by placing one tag on each antenna sequentially (Figure 2E, left). The total PCT increased linearly with the number of tags. The estimated PCT across 24 antennas in the absence of tags was 142 ms (5.9 ms per antenna), and the additional read time per tag was 8.2 ms.

We assessed the effect of the anti-collision algorithm on the PCT by stepwise increasing the number of tags on a single antenna (Figure 2E, right). While this process added an approximately 30 ms delay for the first five tags in both modules, likely due to slot reallocation overhead, its overall impact remained limited.

### 3.2 Behavioral Tracking in Mice using the eeeHive 2D Module

We applied the 2D module to characterize group and individual space use patterns of aged C57BL/6J male mice. By introducing compartmentalization and differential illumination, we generated a spatially heterogeneous environment to facilitate the expression and interpretability of these behaviors (Figures 3A and 3B). Seven-day logs were processed following the workflow in section 3.2.1 (Figure 3C).

#### 3.2.1 Coupling Between Activity Rhythms and Group Dispersion

Activity, defined as the sum of movement distances derived from coordinate changes, was visualized as per-unit-time values for both individuals and the group average (Figures 3D and 3E). As is typical for this strain, mice were primarily active during the dark phase, peaking at both light transitions. The mean daily movement was 253.3 m (SD = 78.7 m), ranging from 149.4 m (M07) to 436.1 m (M01).

To quantify group dispersion, we calculated the mean distance from the centroid (MDC), defined as the average distance of all individuals from the group’s center of mass. MDC fluctuations revealed that the group dispersed during active phases (around 15 cm) and became more cohesive during resting periods (approximately 8 to 9 cm) (Figure 3E).

#### 3.2.2 Spatial Use of the Group

We next quantified the percentage of time spent in each compartment (Rooms 1 to 4 and Center) on an hourly basis for all 24 mice. Throughout the recording, most mice preferred Room 3 due to its red acrylic ceiling that reduced illumination, while a transient Day 1 preference for the similarly shaded Room 4 quickly diminished. Inversely tracking locomotor activity, the time spent in Room 3 decreased during the dark phase, dropping to approximately 70% at dark onset with a smaller decline near the dark-to-light transition (Figure 3F).

Post-experimental visual inspection revealed localized urination marks in the corners of other rooms, with particularly concentrated traces in Room 1 (Supplementary Figure 2A), which was located farthest from Room 3. The amount of bedding material in Room 3 was reduced, but the remaining bedding showed no such accumulation and appeared relatively clean.

These collective spatial-use patterns are consistent with previous findings that mice establish a “home base” with two primary functions: (i) achieving behavioral thermoregulation through group aggregation (49,50), and (ii) the spatial segregation of elimination sites from the primary resting area (51–53).

#### 3.2.3 Spatial Organization of Individuals Within the Shared Home Base

We next examined the spatial organization of individual mice. As an initial approach, we plotted the hourly trajectory of the centroid of spatial positions during the entire experimental period for each individual (Figure 3G). We also calculated the overall centroid across the experimental period, indicated by a red dot in Figure 3G and hereafter referred to as the individual global centroid (IGC). All individuals were primarily stationed in Room 3, confirming that this compartment served as the common home base.

Inter-individual differences were observed in the spatial dispersion of the IGC. Also, IGCs tended to lie closer to the Room 3 cage corner, a more enclosed and potentially secure resting location. These observations prompted us to consider whether the degree of spatial dispersion and the location of the IGC could serve as quantitative descriptors of individual behavioral characteristics.

#### 3.2.4 Inter-Individual Divergence in Roaming Range and Corner Occupancy

To quantify each individual’s spatial dispersion during the experimental period, we calculated the mean distance from the IGC to the animal’s coordinates at all time points. This value was defined as roaming range.

To assess the degree to which each individual’s spatial occupancy was biased toward the Room 3 cage corner, we calculated the mean distance from the corner reference point (coordinate 1,1) to the animal’s position at all time points. This value was defined as distance from the corner (DFC). Day-by-day changes in DFC showed that this value was stable across the experimental period (Supplementary Figure 2B).

A strong positive correlation was observed between roaming range and DFC (r = 0.84) (Figure 3H). Individuals with smaller DFC values exhibited smaller roaming ranges, whereas those farther from the corner showed greater spatial dispersion. These patterns suggested two behavioral tendencies within the group: corner occupiers, whose positions were consistently concentrated near the corner of the home base (e.g., M20, M15, M14, M07), and roamers, whose occupancy extended farther from the corner and who had larger roaming ranges (e.g., M19, M13, M01, M23).

#### 3.2.5 Enhanced Aggregation Near the Home Base Corner

To determine if corner proximity facilitates physical proximity to other individuals, we defined two metrics using the anti-collision capability of the eeeHive: aggregation score (cumulative co-presence time on the same antenna, summed across all co-present individuals) and isolation time (proportion of time without others on the focal or adjacent antennas). DFC correlated negatively with aggregation score (r = -0.80) and positively with isolation time (r = 0.83) (Figures 3I and 3J).

These findings indicate that the Room 3 corner is a highly valuable spatial resource, as it provides a more enclosed environment, distance from latrine sites, and higher local densities that likely reduce thermoregulatory costs. Because this resource is limited, priority access to it may reflect social dominance, distinguishing corner occupier mice from roamers.

#### 3.2.6 Reciprocal Following as Index of Interaction with Other Individuals

We examined whether mice exhibited coordinated movement across individuals. Across the 7-day experiment, the 24 mice made a total of 31,662 room transitions. We analyzed whether these events occurred independently or showed temporal coordination.

First, we defined follow latency as the time elapsed between one individual’s transition from Room X to Room Y and another’s subsequent identical transition. After binning these latencies into 10-second bins, we used a permutation test to determine whether the observed frequencies could arise by chance. Only the 0-10 s bin exhibited a significantly higher frequency than expected, indicating that mice tend to follow another individual within 10 seconds. We defined this action as follow behavior.

We next calculated the ratio of observed to expected events (O/E ratio) using a permutation test to evaluate follow behavior across all 552 directed pairs. O/E ratios ranged from 0 to 9.065 (mean 2.61; maximum Z = 10.5 from M15 to M09). In total, 44.9% of pairs exhibited significantly higher-than-expected follow behavior, and the top 25% of these were visualized as a directed network (Figure 3K)

To examine individual tendencies to follow and be followed irrespective of specific pairs, we correlated out-strength (the sum of Z-scores for following others) and in-strength (the sum of Z-scores for being followed). The resulting moderate positive correlation (r = 0.52; Figure 3L) suggests that individuals are not rigidly divided into leaders and followers but rather vary in their overall engagement in reciprocal following.

Notably, roamers (e.g., M01, M13, M19, M23) showed minimal engagement in reciprocal following, whereas corner occupiers exhibited broader, more diverse interaction profiles. These findings suggest that roamers are more socially independent across both proximity-based metrics and behavioral synchrony.

Correlation analysis of five metrics (DFC, in/out-strength, and in/out-entropy reflecting target uncertainty) revealed that DFC negatively correlated with all other measures, confirming the social independence of roamers (Figure 3M). Furthermore, both strengths positively correlated with their respective entropies, indicating that highly interactive individuals engage with multiple partners with low selectivity.

#### 3.2.7 Multivariate Classification of Individual Behavioral Profiles

To summarize individual behavioral diversity, we performed a principal component analysis (PCA) on 29 standardized metrics. Supplementary Table 2 shows the names and descriptions of each behavioral metric, along with their respective principal component loadings for PC1, PC2, and PC3. The first three principal components (PCs) explained 68.2% of the total variance (Figure 3N). PC1 reflected broad roaming and minimal reciprocal following. PC2 represented high spatial randomness and active following. PC3 captured flattened circadian rhythms with minimal activity differences between light and dark phases. Supplementary Figure 2C presents the pairwise biplots of the three principal components.

Subsequent hierarchical clustering of these PC scores using Ward’s method identified four profiles (Figures 3O and 3P). Cluster 1 (n = 3, high PC1) corresponded to roamers, while Clusters 2 (n = 9) and 3 (n = 6) were characterized by high PC2 and PC3 scores, respectively. Cluster 4 (n = 6) exhibited low scores across all PCs and primarily consisted of corner occupiers.

### 3.3 Behavioral Tracking in Marmosets Using the eeeHive FLEX Module

To demonstrate the FLEX module’s applicability, we quantified individual behaviors and inter-individual interactions within a marmoset colony (Figures 4A and 4B). For further validation, we confirmed that the eeeHive data and our interpretations were consistent with independent daily observational records from animal caretakers (Supplementary Information 3).

To validate data acquisition, we compared system detections against video recordings over 15 days (09:00–15:00). All 850 access events detected by the antennas completely matched the video observations, confirming tracking accuracy for group-housed marmosets. For the main analysis, seven weeks of access logs were recorded and processed as described in Materials and Methods 3.3.2.2.

#### 3.3.1 Long-Term Behavioral Log Visualization

Behavioral log visualizations (Figure 4C; Supplementary Animation 2) highlighted frequent daytime visits to feeder and water stations across all marmosets yet demonstrated consistent inter-individual variation in prolonged nestbox occupancy during the dark phase.

Over the 7-week observation period, F1, M1, M3, and M4 spent over 90% of the dark phase inside the nestbox (see Figure 4I for specific nestbox occupancy). In contrast, F2 and M2 spent only 54.9% and 74.4% of their time there, respectively. Instead, both individuals lingered substantially at the entrance antennas (28.3% [166.3 h] and 22.1% [129.7 h]), with F2 spending an additional 16.8% (98.7 h) entirely outside.

#### 3.3.2 Transition Probability Mapping Between Locations

To characterize individual movement patterns, we applied a first-order Markov chain model treating each antenna location (Nest 1–6, Feeder 1–2, Water station 1–2) as a state. The resulting transition probabilities were visualized as individual network graphs (Figure 4D) and used to calculate transition entropy. F2 exhibited the lowest entropy (strongly biased movements, peaking at an 11% transition from Feeder 1 to 2), whereas M4 showed the highest entropy (highly unpredictable transitions, peaking at 4%).

#### 3.3.3 Clustered Nestbox Visits During the Light Phase

During the light phase, marmosets mostly remained outside but frequently exhibited “clustered visits” (multiple individuals entering and exiting the nestbox together). The duration of these visits, measured from the simultaneous entry into an empty nestbox until complete evacuation, was typically under 300 seconds (Figure 4E). This temporal clustering suggests that marmosets engage in coordinated follow behavior during daytime nestbox use.

#### 3.3.4 Operational Definition and Classification of Follow-Related Behaviors

Based on daytime clustered visits, we operationally defined four follow-related events between individuals A and B at a given nest: (i) following entry (B enters within 60 s of A’s entry), (ii) avoidance (A exits within 60 s of B’s entry), and (iii) following exit (B exits within 60 s of A’s exit). A sequential combination of these three events was defined as (iv) chasing. We adopted this 60-s threshold because latency distributions revealed that most events occurred within this interval (Supplementary Figure 3).

#### 3.3.5 Pair-Specific Strength of Follow Relationships

To quantify pair-specific follow relationships while controlling for chance encounters, we compared the observed combined follow events (entries plus exits) to expected values derived from 1,000 permutations. By applying a 60-minute block circular shift to one individual’s light phase data, we randomized temporal alignment while preserving intrinsic movement patterns.

Across all 30 directed relationships (15 pairs), the O/E ratio was significantly greater than chance (p < 0.05, Figure 4F), averaging 12.5 and ranging from 4.79 (M2 following M3; observed = 51, expected = 10.7) to 24.3 (M3 following F1; observed = 561, expected = 23.1). Thus, consistent yet highly variable follow behavior existed throughout the group. Notably, F1 was intensely followed by M3 and M1 (O/E ratios of 24.3 and 23.1, respectively), while F2 was strongly followed by M1 and her same-sex peer F1 (O/E ratios of 19.5 and 16.5).

We next analyzed chasing, a more intensive follow behavior interpreted as the persistent pursuit of an escaping individual. Across the 30 directed relationships, the O/E ratio for chasing averaged 17.8 and ranged from 1.55 (M2 chasing M3; observed = 1, expected = 0.646) to 56.1 (M3 chasing F1; observed = 72, expected = 1.284; Figure 4G). All ratios except the 1.55 minimum were statistically significant (p < 0.05). These findings confirm that chasing occurs with even greater pair-to-pair variability than combined follow events. Notably, F1 was persistently chased by M3 and M1 (O/E ratios of 56.1 and 34.6, respectively), while F2 was intensely pursued by M1 and her same-sex peer F1 (O/E ratios of 39.2 and 40.6).

#### 3.3.6 Divergence in the Strength and Asymmetry of Follow Interactions

We next examined the relationship between the volume and asymmetry of follow behaviors. Defining bidirectional follow strength and follow asymmetry as the sum and difference of the paired Z-scores, respectively, we found a moderate positive correlation between the two metrics (r = 0.44, Figure 4H).

The three most asymmetric pairs (M3 and M2, M3 and F1, M1 and F1) aligned perfectly with daily caretaker observations (Supplementary Information 3). Specifically, the M3 and M2 asymmetry reflected clear dominance, with M2 emitting submissive vocalizations while retreating from M3’s threats and chases. The M3 and F1 asymmetry corresponded to repeated mating behaviors and persistent pursuit by M3. Finally, although M1’s overall low activity likely obscured visually obvious following, caretakers independently characterized the M1 and F1 relationship as highly affiliative.

#### 3.3.7 Individual Differences in Dark Phase Nestbox Selection

As nocturnal nestbox sharing may provide insights into social relationships, we mapped the primary nighttime locations of all individuals over seven weeks (Figure 4I). Overall, all individuals relied on Nest 1 as their default base, making occasional excursions to other locations. While M4 remained almost exclusively in Nest 1 throughout the experiment, other individuals displayed distinct spatial dynamics during these relocations. Specifically, F1 and M3 often co-occupied Nest 3. Furthermore, on the three occasions M1 left Nest 1, he always stayed with F1 in Nest 3. Additionally, M2 and F2 exhibited solitary tendencies mid-experiment, with M2 increasingly resting alone in Nest 3 or 4 from Day 20, and F2 spending the night completely outside any nestbox from Day 19.

#### 3.3.8 Pairwise Nestbox Co-Occupancy During the Dark Phase

To visualize nighttime spatial associations, we mapped the total dark phase nestbox co-occupancy for each pair (Figure 4J). Co-occupancy averaged 441.3 hours, ranging from 319.3 hours (M2 and M3) to 547.5 hours (M1 and M4). Consistent with their highly asymmetric daytime pursuit of F1, both M1 and M3 shared long nocturnal co-occupancy with her. In contrast, the strongly asymmetric same-sex M3 and M2 pair showed the lowest co-occupancy time, highlighting a clear dissociation between daytime follow dynamics and nighttime spatial affiliation.

#### 3.3.9 Coupling Between Nighttime Nestbox Co-Occupancy and Daytime Follow Dynamics

To examine whether daytime follow behavior and nighttime nestbox co-occupancy were interrelated, we plotted each individual’s total bidirectional follow behavior (sum of Z-scores for combined follow events) against their mean dark-phase co-occupancy time. The scatter plot revealed a strong positive correlation (r = 0.94, Figure 4K). Notably, M2 was clearly isolated with low values on both axes, indicating a persistent tendency to remain apart from the group across both active and resting phases.

### 3.4 Application of eeeHive in Aquatic Environments

While Figure 2D demonstrates that water minimally affects tag detection range, we conducted experiments to confirm the system’s practical applicability for analyzing group-housed newts and tilapia in freshwater.

#### 3.4.1 Spatial Organization and Cohesion in Newts

Validating the 2D module’s underwater accuracy, system-detected coordinates for newts showed complete agreement with manually annotated positions across 68 video frames sampled at 10-minute intervals.

Behavioral analyses utilized 136 h (∼5.5 consecutive days) of log data, excluding a 6-h facility-related interruption on the third day. Data processing followed the procedures detailed in Materials and Methods 3.3.2.1.

##### 3.4.1.1 Individual Variability in Activity Levels in Newts

Quantifying activity levels (travel distance) revealed comparable baselines for F1, F2, and M1 (Figure 5B). In contrast, M2 exhibited an approximately fourfold higher value throughout the recording period (Supplementary Figure 4A).

##### 3.4.1.2 Peripheral Spatial Bias in Newts

Analysis of spatial utilization revealed a universal preference for peripheral regions (tank walls and corners), accounting for most of the dwelling time (Figure 5C) and approximately 70% of all transitions (Supplementary Figure 4B). Within this strong peripheral bias, individual spatial strategies diverged (Figure 5D). Females (F1, F2) utilized the central area more frequently than males. Meanwhile, the males exhibited distinct peripheral preferences, with M1 residing primarily in corners and M2 along the walls.

##### 3.4.1.3 Transient Group-Level Cohesion in Newts

To evaluate group dispersion, we calculated the mean distance from centroid (MDC) per second. The animals exhibited recurrent, transient cohesion episodes (aggregation within a few centimeters) that were independent of feeding (Supplementary Figure 4C). To determine if these events occurred by chance, we compared the observed MDC distribution against 1,000 permutations using a 10-minute block circular shift. MDC between 0 and 90 mm showed significantly elevated O/E ratios (p < 0.05), with the ratios increasing progressively in smaller MDC bins (Figure 5E). This indicates that the newts tend to form tight clusters more frequently than expected by chance.

##### 3.4.1.4 Pair-Specific Proximity Structure in Newts

Defining proximity as the cumulative time (s) spent within ≤ 100 mm of each other, we assessed all six pairs via 1,000 permutations. Every pair except the M1 and M2 dyad spent significantly more time in proximity than chance expectations (Figure 5F). The strongest association occurred between F2 and M2 (O/E = 1.39, observed = 92,375 s, expected = 66,632 s; p < 0.001), contrasting with the non-significant male and male pair (O/E = 0.98, observed = 65,942 s, expected = 67,033 s).

#### 3.4.2 Compartment Transitions and Group Structure in Tilapia

To validate the FLEX module’s accuracy under fully submerged conditions (Figure 5G), we compared 30 min of system data against simultaneous video recordings. After applying our chattering-removal and transition-definition rules, the system achieved 100% agreement with video observations across all 207 bidirectional compartment transitions. Subsequent behavioral analyses utilized 75.5 h of continuous recording data, processed as detailed in Materials and Methods 3.3.2.2.

##### 3.4.2.1 Light–Dark–Entrained Activity Rhythms in Tilapia

Quantifying activity as the number of inter-compartment transitions, we observed a clear diurnal pattern across all tilapia, featuring increased movement during the light phase (Figure 5H). Total transitions varied individually over the experimental period, averaging 1,019 per fish and ranging from 623 (M4) to 1,278 (M6) (Supplementary Figure 4D).

##### 3.4.2.2 Pairwise Co-Presence in Tilapia

Visualizing compartment occupancy suggested temporal synchrony in transitions among individuals (Figure 5I). To quantify this, we defined co-presence (%) as the proportion of time each of the 55 pairs shared a compartment. We then compared observed values against 1,000 permutations using a 10-min block circular shift. The empirical co-presence distribution shifted markedly higher than the null expectation (Figure 5J), indicating that compartment use was not independent, with co-presence occurring more frequently than expected by chance.

While 44 of the 55 pairs demonstrated significantly greater co-presence than chance expectations (Supplementary Figure 4E), the overall effect sizes were modest, reaching a maximum O/E ratio of only 1.11 between M5 and M11 (observed = 0.64, expected = 0.58).

##### 3.4.2.3 Following Tendency in a Group of Tilapia

To determine if compartment transitions were temporally associated, we compared the latencies of consecutive same-direction movements between individuals against a null distribution generated via 1,000 permutations (10-min block circular shifts). Short following latencies (< 4 s) occurred significantly more frequently than expected by chance (p < 0.05, Figure 5K). This synchronized timing demonstrates that tilapia have tendency to immediately follow one another, reflecting the rapid propagation characteristic of fish shoaling behavior.

##### 3.4.2.4 Pairwise Follow Structure in Tilapia

At the pair level, we evaluated short-latency follow events (< 3 s, derived from the group analysis). Permutation testing (1,000 single-individual 10-min block circular shifts) revealed that 62 of 110 dyads exhibited significantly higher follow counts than expected. With effects peaking from M11 to M7 (O/E = 2.16) and frequent directional asymmetries, these interactions constitute a directed network (Figure 5L).

##### 3.4.2.5 Structural Organization of Reciprocal Follow in Tilapia

To characterize reciprocal following, we defined out-strength and in-strength as the sum of Z-scores for follow interactions directed toward and received from other group members, respectively. Plotting these measures (Figure 5M) revealed heterogeneous individual engagement (e.g., active M7 vs. inactive M8) with a non-significant moderate correlation (r = 0.43). Notably, many individuals exhibited asymmetrical profiles, acting primarily as either followers or leaders. This suggests that tilapia collective movement emerges from diverse interactive roles rather than a simple linear hierarchy.

## 4 Discussion

Pioneered by Ely et al. (54), integrating automated measurement techniques with species-specific enclosures is a powerful approach for ethological research. RFID serves as a core technology here, enabling flexible and scalable implementation of antennas for precise individual tracking within social groups (23,30,31,55).

To evaluate the scientific validity and feasibility of HF RFID in this context, we developed the eeeHive system. Below, we discuss the advantages and concerns regarding HF RFID, summarize our experimental findings, and explore its broader implications across multiple fields, ultimately advocating for its standardization.

### 4.1 From LF to HF: Rationale and Dispelling Concerns

#### 4.1.1 Technical Constraints of Conventional LF RFID Systems

Restricted by low data rates, LF RFID’s slow polling forces a severe spatiotemporal trade-off: denser antennas risk missing swift animal movements. Conversely, the high data rate of the HF RFID-based eeeHive (5.9 ms/antenna polling, 8.2 ms/tag reading) permits a much higher antenna density while maintaining rapid polling cycles, ultimately achieving high-resolution behavioral tracking in both space and time.

The second major limitation of LF RFID is severe data loss from tag collisions when multiple animals share an antenna field. This issue is critical for densely aggregating species like mice. Indeed, our data revealed that individuals were co-located on the same antenna 48.7% of the time (56.4% light, 41.0% dark phase). These high aggregation rates underscore that simultaneous multi-tag detection, a core feature of HF RFID, is indispensable for accurate tracking in group-housed conditions.

#### 4.1.2 Historical Context of LF RFID Standardization in Animals

The technological foundation for passive inductive animal tags was established through a series of patents by the end of 1970s, which were contemporaneously applied to livestock management (56,57). In the 1980s, the growing demand for injectable and implantable tags drove the refinement of circuit miniaturization and encapsulation technologies (58,59). This technological innovation was strongly propelled by national demands in fisheries management and ecological monitoring, notably the large-scale tracking of salmonid migration in the Columbia River Basin (60,61). Consequently, early implantable animal tags were developed with a strong emphasis on underwater use, establishing the adoption of the LF band, which is less susceptible to signal attenuation by conductive media such as water and body tissues, as a physical prerequisite.

Subsequently, through the 1990s, LF tags were increasingly adopted for the management of companion animals, as well as for the observational studies of various wild and captive animals (23,62). In the context of laboratory animals, their safety for long-term implantation and utility for individual identification were validated in mice, rats, woodchucks, rabbits, and amphibians (20,63,64).

However, during this decade, individual companies designed systems based on proprietary frequencies (e.g., 400 kHz, 125 kHz, and 128 kHz) and specifications, resulting in the lack of compatibility among products. To resolve this issue and foster a global market, the ISO initiated the standardization of RFID systems for animal identification in the mid-1990s. This effort culminated in the publication of ISO 11784 (40,65) and ISO 11785 (39), which established 134.2 kHz as the global standard operating frequency.

Meanwhile, HF RFID systems, particularly those based on ISO/IEC 14443 and ISO/IEC 15693 standards, achieved widespread global adoption in the early 2000s following the establishment of international standards and the expansion of contactless smart card applications (66). By that time, the animal-related applications had already established LF-based equipment, supply chains, field operations, and database systems built around ISO 11784/11785. Transitioning to HF would have required a large-scale infrastructural overhaul, for which there was likely no economic incentive.

Consequently, the continued dominance of LF RFID in animal identification and behavioral tracking to this day can be attributed to two main factors: its physical characteristic of being less susceptible to interference from conductive media (such as water and body tissues), and the powerful historical inertia generated by the early proliferation and standardization of LF-based infrastructure.

#### 4.1.3 Limited Adoption of HF RFID in Behavioral Research

HF RFID has nevertheless been applied to behavioral tracking in specific research contexts, particularly in studies of social insects such as bees and ants, where high-throughput tracking of large numbers of individuals is required (67–69). This has been enabled by the availability of miniaturized HF RFID tags (e.g., mic3-tag, ∼1.5 mm³; Microsensys GmbH, Erfurt, Germany) that can be attached to the body surface of insects, which can only be activated at very short distances (≤ 5 mm) from the antenna.

A notable example in vertebrates further illustrates the advantages of HF RFID: a study on Northern carmine bee-eaters conducted in a zoo showed that replacing an LF system with an HF system improved data acquisition, enabled simultaneous detection of multiple individuals, and reduced missed detections in nest visitation records (70). Although a few other ethological studies have utilized HF RFID, their applications have remained largely restricted to specific research areas and a limited number of species, primarily in field settings (71).

To the best of our knowledge, no AHCM system leveraging the advantages of HF RFID has yet been practically implemented for a variety of laboratory animals such as rodents. However, given the growing demand for ethological approaches in the laboratory, HF RFID should be widely adopted as a standard laboratory technique.

#### 4.1.4 Addressing Common Concerns in the Adoption of HF RFID

In this section, we outline two potential concerns associated with adopting HF RFID as a standard method in animal experiments and explain why these do not constitute substantial barriers to its implementation.

##### 4.1.4.1 Effects of Conductive Media on RFID Tag Reading Performance

While LF RFID is widely favored for its low signal attenuation in conductive media, our data demonstrate that HF RFID is equally robust for the short-range communication typical of AHCM. This can be quantitatively understood using penetration depth (the distance at which the electromagnetic field decays to 1/e). According to Benelli and Pozzebon (2013), the HF penetration depth is 30–2,500 cm in freshwater and 6.8 cm in highly conductive seawater (72). Because AHCM typically utilizes within a ∼40 mm tag–antenna distance, this theoretical decay is negligible. Consequently, reliable tag detection remains entirely feasible whether passing through biological tissues, freshwater, or seawater.

##### 4.1.4.2 Effects of Exposure to HF Electromagnetic Fields on Animal Health

To date, no established adverse health effects have been identified from long-term exposure to electromagnetic fields in the HF frequency range. Long-term evaluations by international expert bodies such as the International Commission on Non-Ionizing Radiation Protection (ICNIRP) and the World Health Organization (WHO) have consistently concluded that exposure to radiofrequency electromagnetic fields below guideline limits is not associated with adverse health effects. HF-band RFID systems operate well within these limits (73,74).

As expected, no adverse health effects associated with HF tag implantation were observed over years in all animal species used in the present study (Supplementary information 4). Yet, continued attention should be given to the potential long-term health effects of exposure to alternating magnetic fields associated with HF RFID.

### 4.2 Key Findings and Practical Considerations in eeeHive Identified through Experiments

#### 4.2.1 Mice Experiment

The primary objective of this experiment was to evaluate the functionality and scalability of the eeeHive 2D module. We confirmed a single host computer could handle simultaneous data acquisition from four 2D modules (96 antennas in total) within the polling cycle time equivalent to a single 2D module (24 antennas). The computational load on the host computer remained negligible. Furthermore, time-synchronized logs were successfully collected, integrated, and analyzed as a single dataset.

Behavioral analyses revealed two major findings. First, all 24 individuals rapidly aggregated to establish a single shared home base (Room 3) within the first day, spending most of the subsequent 7 days there. Bedding in this compartment showed minimal evidence of urination, demonstrating a strict collective segregation of resting and elimination sites. These behaviors perfectly align with previous reports on murine behavioral thermoregulation through aggregation (49,50), and their strong preference for latrine segregation (51–53). This rapid and highly reproducible coordination among a relatively large group appears to suggest that mice possess a greater capacity for collective behavior than typically recognized.

Second, despite this common tendency, individuals displayed diversity in spatial usage, proximity to conspecifics, follow behavior, and diurnal activity patterns. Notably, two distinct behavioral characteristics stood out within the home base: individuals that consistently occupied corner areas (corner occupiers) and those stayed away from corners (roamers).

These characteristics may reflect a social dominance competing for a limited spatial resource as a comfortable spot furthest from elimination sites and enclosed by walls. Consistent with this interpretation, previous studies in inbred mice have shown that subordinate individuals tend to be displaced toward areas near elimination sites, linking spatial positioning to social status (54,75). Corner occupiers formed dense clusters in close proximity to multiple conspecifics, whereas roamers occupied regions with lower local density and showed reduced temporal proximity during inter-compartment transitions, suggesting a tendency toward social isolation.

The spatial partitioning of the living environment facilitated the interpretation of these results. Further efforts to compartmentalize environments with heterogeneous conditions, such as those exemplified by the visible burrow system and Eco-HAB, are expected to reveal a broader range of behavioral characteristics (55,76–78).

#### 4.2.2 Marmosets Experiment

The primary objective of this experiment was to evaluate the applicability of the eeeHive FLEX module for tracking freely moving marmosets in a large enclosure. In this setup, multiple ring-type antennas were deployed across the experimental space. The cables connecting the reader and antennas ranged from approximately 2 to 10 m in length, and all operated without troubles.

A key consideration in this setup is the pair of antennas installed in each nestbox entrance tube, used to infer movement direction from the temporal difference in tag detection. To ensure reliable detection of this time difference, the tube diameter was adjusted to induce deceleration of marmosets during passage. As noted in previous studies using LF RFID systems (31,32), structural modifications of the tube, such as narrowing or bending sections, can improve detection reliability within ring antennas. Importantly, such design considerations remain equally important in HF RFID-based systems.

This type of experimental setup is not intended to capture full spatial trajectories but is instead well suited for logging interpretable behavioral events, such as nestbox entry/exit and access to food or water. It should be noted that similar experimental designs can also be implemented using LF RFID-based multi-antenna systems, provided that an independent-readers configuration is adopted (i.e., without sequential polling), although such systems lack anti-collision capability. In the present marmoset data, the proportion of simultaneous detection of two or more tag IDs ranged from 0.1–5.6% across individuals over the entire recording period, suggesting that tag collisions are not negligible.

The eeeHive FLEX module successfully characterized behavioral features and inferred social relationships within a marmoset colony. Temporal proximity and sequential order of nestbox entry and exit during the daytime, as well as patterns of nestbox co-occupancy at night, served as informative indicators of social relationships among individuals. These results were highly consistent with independent visual observations by animal caretakers during the daytime.

For example, eeeHive revealed that M3 exhibited pronounced chasing behavior toward F1 (Figure 4G). This was consistent with the caretakers’ observations that M3 persistently followed F1 and engaged in mating behavior. F1 subsequently became pregnant and gave birth, suggesting that such chasing reflected the formation of a reproductive pair between M3 and F1.

In contrast, chasing behavior between same-sex individuals was interpreted as a dominance-related interaction. The chasing of F2 by F1 revealed by the eeeHive data was consistent with their hierarchical relationship, with F1 exhibiting behaviors characteristic of a dominant individual (79), unequivocally recognized by caretakers as the dominant, while F2 was subordinate. Similarly, the unidirectional interaction in which M3 consistently chased M2 (Figure 4H) aligned with caretaker observations, which described persistent and aggressive chasing by M3 accompanied by submissive behaviors in M2 (79), including frequent infant-like vocalizations. These findings are consistent with the social ecology of marmosets, in which breeding individuals (F1 and M3) form a female-led or pair-based dual dominance hierarchy (80–82).

The individuals identified as subordinate (F2 and M2) exhibited distinctive use of the nestbox during nighttime (Figures 4C and 4I). Although they typically slept in Nest 1, they were often located in the narrow entrance tube rather than inside the nestbox, suggesting exclusion from the core aggregation. From the middle of the experiment, M2 increasingly slept alone in a different nestbox, while F2 was more frequently found outside the nestbox. These behaviors are consistent with early stages of social exclusion in marmosets, which are characterized by increased submissive vocalizations and disrupted co-sleeping (83). Supporting this, M2 frequently showed submissive vocalizations toward M3 and was later subjected to aggression. F2 was ultimately excluded from the group approximately one year after the experiment.

Notably, the phenomena observed in these two individuals share key features with the “roamer” characteristic in mice experiment, in terms of unstable and restricted access to the nestbox as a limited spatial resource, as well as reduced social interactions with other individuals. This similarity raises the possibility that such behavioral patterns may represent a generalizable phenomenon across species. A more detailed analysis of spatial positioning within the nestbox may enable a more precise characterization of their social relationships.

#### 4.2.3 Newts and Tilapia Experiment

The aim of this study was to validate that the eeeHive 2D and FLEX modules can operate reliably in freshwater environments and enable continuous tracking of multiple individuals within social groups. In conclusion, evaluation of tag detection distance confirmed that detection distance in freshwater was comparable to that in air, indicating that the system is practically applicable to freshwater organisms.

Consistent with studies in terrestrial environments using mice and marmosets, eeeHive enabled the analysis of group dispersion, individual activity and spatial utilization, as well as pairwise co-presence and follow relationships in group-housed newts and tilapia under aquatic conditions. While the small sample size and short tracking period preclude a detailed discussion of the individual data, it was evident that newts tend to exhibit inter-individual cohesion, and tilapia show a propensity for co-presence within the same compartment and synchronized movements between compartments.

As in terrestrial experiments, introducing more complex spatial partitioning of the environment may further extend the applicability of the eeeHive-based approach to investigate a wider range of phenomena in aquatic ethology.

### 4.3 Implications for Various Research Fields

#### 4.3.1 Implications for Ethology-driven Biomedical Research

Recent ethological research on small laboratory animals, including rodents and fish, has increasingly revealed cognitive capacities that are not readily explained by simple stimulus–response (S–R) models (84,85). A notable example is the discovery of intentional tactical deception called “deceptive dodging” in wild black-striped mice (86), which involves a sequence of strategic escape actions. Specifically, evading the pursuer’s field of view, remaining motionless and silent to conceal its presence, and subsequently escaping when the pursuer’s attention is diverted. It likely requires cognitive functions such as intention understanding, future state prediction, and inhibitory behavioral control.

However, such behavioral complexity may not be entirely unexpected. Many rodent species, likely originating shortly after the Cretaceous–Paleogene boundary (∼62 million years ago), have persisted without substantial morphological anti-predator defenses (87). From the perspective of the predatory intelligence hypothesis (88), which posits that the cognitive challenges of predator evasion drive an evolutionary arms race, these observations are consistent with an adaptive framework in which even small animal models with simple brain structures may possess cognitive capacities even partially relevant to human brain functions.

If such behaviors like deceptive dodging can be efficiently reproduced on a larger scale within controlled laboratory environments using systems like the combination of eeeHive 2D and FLEX modules, it would become possible to systematically investigate the genetic and environmental factors, as well as the effects of pharmacological interventions, that govern their expression. Uncovering the underlying mechanisms of these cognitive behaviors could accelerate translational research aimed at overcoming the cognitive deficits associated with a wide spectrum of neurological and neuropsychiatric disorders.

#### 4.3.2 Implications for Drug Efficacy and Safety Assessment

By enabling continuous monitoring of spontaneous behaviors within minimally disturbed living environments, AHCM provides a framework for time-resolved evaluation of drug effects and reduces the risk of overlooking unexpected behavioral or physiological changes (89). The scalability and flexibility of HF RFID–based systems such as eeeHive can further expand the scope of AHCM applications. High spatiotemporal resolution tracking across diverse enclosure designs enables more multidimensional assessment of drug efficacy, safety, and toxicity. In addition, quantification of inter-individual distance, aggregation, and movement synchrony within groups may provide particularly informative metrics relevant to neuropsychiatric domains. The ability to apply comparable metrics across multiple animal species also facilitates evaluation of translational relevance.

The capabilities of eeeHive are well aligned with approaches that leverage large-scale behavioral datasets collected under standardized conditions to classify compounds across multiple pharmacological categories and to screen novel agents (90). These features are also particularly valuable in environmental toxicology, where the vast number of substances and the difficulty in predicting both the timing and endpoints of toxicity underscore the need for scalable and unbiased behavioral screening frameworks.

#### 4.3.3 Implications for Animal-centric Refinement

In the care and management of laboratory animals, applying refinement from an animal-centric perspective is a major challenge (91). A promising approach is to define what environmental conditions animals prefer or avoid using AHCM. For instance, RFID-assisted preference choice tests have successfully identified preferred nesting materials and structural elements and so on, with ongoing progress in optimizing evaluation metrics and analyses (92,93).

We also emphasize the importance of analyzing precise individual locations, movement trajectories, and inter-individual relationships for identifying aversive environmental factors within the cage. The present study demonstrated the strong aversive effect of the latrine areas in mice and the inequity in nocturnal nestbox access for subordinate marmosets. These findings have provided us inspirations for resolving problems in animal housing environments. We anticipate that the broader adoption of systems like eeeHive will pave the way for major contributions to the refinement of future animal experiments.

## 6 Author Contributions

T.E. and S.B. conceived and designed the study. S.B., S.S., and T.E. conducted the experiments and performed the analyses. T.E., K.H., and T.K. developed and provided analytical tools. T.E., S.B., H.Y., and H.P.L. contributed to experimental design and data interpretation. T.E. and S.B. drafted the manuscript, and all authors contributed to its revision for important intellectual content.

## Acknowledgments

We sincerely thank Kayoko Taki (Phenovance LLC) for her valuable assistance with experiments and laboratory maintenance. Sincere gratitude is extended to Hitoshi Akimoto (Beyonstem Inc.) and Naoki Yaegashi (Fluorite Technologies Inc.) for their invaluable support in the fabrication, design, and functional evaluation of eeeHive. Special thanks to Rie Gonda, Miyuki Suzuki, and Emiko Hatano from the Department of Psychiatry at Hamamatsu University School of Medicine for their extraordinary contributions to the daily care of marmosets and their technical assistance. We are grateful to Dr. Hiroaki Kogure (Kogure Consulting Engineers, Tokyo, Japan) for providing a model of the magnetic field strength distribution, and to Dr. Takahiro Ogawa (MEL Inc., Nagoya, Japan) for providing access to the S-NAP® Wireless Suite software used in this research. We thank Synics AG (Regensdorf, Switzerland) for technical support. We also thank Prof. Frank Rühli for scientific help and Irina Lipp (Institute of Evolutionary Medicine University of Zürich) for valuable administrative support.

## Supplementary Material

Supplementary Animation 1. Tracking of group-housed mice using eeeHive 2D.

Supplementary Animation 2. Tracking of group-housed marmosets using eeeHive FLEX.

Supplementary Table 1. Configurations for Multi-Antenna RFID Systems

Supplementary Table 2. Behavioral Metrics and Principal Component Loadings

Supplementary Figure 1. Supplementary Figures for the Tag Detection Range

Supplementary Figure 2. Supplementary Figures for the Mice Experiment.

Supplementary Figure 3. Supplementary Figures for the Marmosets Experiment.

Supplementary Figure 4. Supplementary Figures for the Aquatic Environment Experiment.

Supplementary Information 1. Supplementary Methods for Evaluation of Tag Read Range and Processing Speeds

Supplementary Information 2. Marmoset diet

Supplementary Information 3. Observational Report by the Marmoset Caretakers Supplementary Information 4. Safety evaluation of HF RFID tag implantation

## 8 Data Availability Statement

The datasets analyzed for this study are available from the corresponding author upon reasonable request.

