## Supplementary Material for "eeeHive: a new HF RFID-based automated behavioral monitoring system for group-housed animals with high spatiotemporal resolution"

**Supplementary Animation 1. Tracking of group-housed mice using eeeHive 2D.**

Time-lapse visualization of positional detections for all mice over the 7-day experimental period, based on antenna-derived spatial data. Dark background indicates dark period: lights on at 8:00, lights off at 20:00.

**Supplementary Animation 2. Tracking of group-housed marmosets using eeeHive FLEX.**

Time-lapse visualization of antenna-access events in common marmosets. Recordings from Week 1, Week 2, and Week 5 are shown to illustrate temporal changes in access patterns. Dark background indicates dark period: lights on at 7:00, lights off at 19:00. NEST, FOOD, etc. indicate antenna locations (multiple nest boxes, feeders, water stations). The videos are trimmed during the dark period because they rarely move at night.

### Supplementary Table 1. Configurations for Multi-Antenna RFID Systems

| Configuration type | Sequential polling | Structural features | Advantages | Constraints | System examples |
|---|---|---|---|---|---|
| Independent-readers configuration | No | Deploys multiple independent reader-antenna units at distances sufficient to prevent mutual interference. | Allows all antennas to remain continuously active, eliminating the risk of missed tag reads caused by temporal antenna deactivation for sequential polling. | Requires low antenna density to prevent interference, resulting in low spatial resolution.<br>Still suffers from data rate-dependent time constraints. | Lipp et al., 2013;<br>Konig et al., 2015;<br>Puscian et al., 2016 |
| Reader-switching configuration | Yes | Deploys multiple reader–antenna units, activated sequentially via a master controller to prevent mutual interference. | Enables denser antenna placement, resulting in high spatial resolution. | Introduces temporal blind spots (antenna deactivation) due to sequential polling, which are exacerbated by low data rates.<br>Incurs complicated and space-consuming architecture. | de Chaumont et al., 2019;<br>Habedank et al., 2022 |
| Antenna-multiplexing configuration | Yes | Deploys multiple antennas connected to a single reader via a multiplexer, activating them sequentially to prevent mutual interference. | Enables denser antenna placement, resulting in high spatial resolution. Offers a simple and space-efficient architecture. | Introduces temporal blind spots (antenna deactivation) due to sequential polling, which are exacerbated by low data rates. | Dell'Omo 1998, 2000;<br>Redfern et al., 2017;<br>eeeHive in the present study |

### Supplementary Table 2. Behavioral Metrics and Principal Component Loadings

| Index | Description | PC1 | PC2 | PC3 |
|---|---|---|---|---|
| TDL | Total movement distance during the light phase | 0.380773 | -0.4092 | 0.679604 |
| TDD | Total movement distance during the dark phase | 0.774027 | -0.4666 | 0.122741 |
| TDR | Ratio of TDL to TDD | 0.539372 | 0.006066 | -0.77409 |
| AMP | Amplitude of circadian activity rhythm derived from cosinor analysis | 0.589177 | -0.40945 | 0.282577 |
| ACP | Acrophase of circadian activity rhythm derived from cosinor analysis | 0.646518 | -0.3466 | -0.55757 |
| OFF2 | Ratio of activity during the two hours surrounding lights off | -0.12932 | 0.469534 | -0.60986 |
| ON2 | Ratio of activity during the two hours surrounding lights on | -0.32652 | -0.18333 | 0.810366 |
| Ratio2 | Ratio of OFF2 to ON2 | 0.049865 | 0.388074 | -0.80449 |
| STP_L | Utilization rate of the home base (room 3) during the light phase | -0.30363 | -0.48717 | -0.68656 |
| STP_D | Utilization rate of the home base (room 3) during the dark phase | -0.92599 | -0.04186 | -0.02989 |
| DFO_L | Mean distance from the corner of the home base (room 3) during the light phase | 0.770084 | 0.336384 | 0.337633 |
| DFO_D | Mean distance from the corner of the home base (room 3) during the dark phase | 0.945987 | 0.06939 | -0.03383 |
| RR | Hourly variance of the spatial centroid | 0.778358 | 0.236279 | 0.312943 |
| AANE_L | Antenna access entropy during the light phase | -0.08636 | 0.739893 | 0.165879 |
| AANE_D | Antenna access entropy during the dark phase | 0.232534 | 0.760902 | -0.21385 |
| ASTE_L | Antenna dwell time entropy during the light phase | 0.57583 | 0.594624 | 0.489628 |
| ASTE_D | Antenna dwell time entropy during the dark phase | 0.893948 | 0.313458 | -0.01617 |
| RMP_3 | Most frequent sequence of three consecutive locations (e.g., Room 3 to Room 2 to Room 3) | 0.693121 | -0.48478 | 0.17577 |
| RMP_3E | Entropy of the area transition sequence of three consecutive locations | -0.27761 | 0.718808 | -0.32209 |
| TM_L | Proportion of movement along the walls during the light phase | 0.206946 | -0.30259 | -0.34719 |
| TM_D | Proportion of movement along the walls during the dark phase | -0.57249 | -0.0873 | 0.487956 |
| AGGE_L | Entropy of aggregation partners during the light phase | 0.307243 | 0.473504 | 0.204919 |
| AGGE_D | Entropy of aggregation partners during the dark phase | 0.512061 | 0.32428 | -0.29896 |
| AGG_L | Aggregation score during the light phase | -0.60218 | -0.50723 | -0.31499 |
| AGG_D | Aggregation score during the dark phase | -0.86994 | -0.27437 | 0.097348 |
| Follow_Out_Strength | Propensity to follow others (Sum of Z) | -0.4633 | 0.561695 | 0.060243 |
| Follow_In_Strength | Propensity to be followed by others (Sum of Z) | -0.44595 | 0.437255 | 0.23164 |
| Follow_Out_Entropy | Entropy of followed individuals | -0.30103 | 0.62527 | 0.296814 |
| Follow_In_Entropy | Entropy of followers | -0.50575 | 0.265793 | 0.162009 |

**Supplementary Figure 1.**

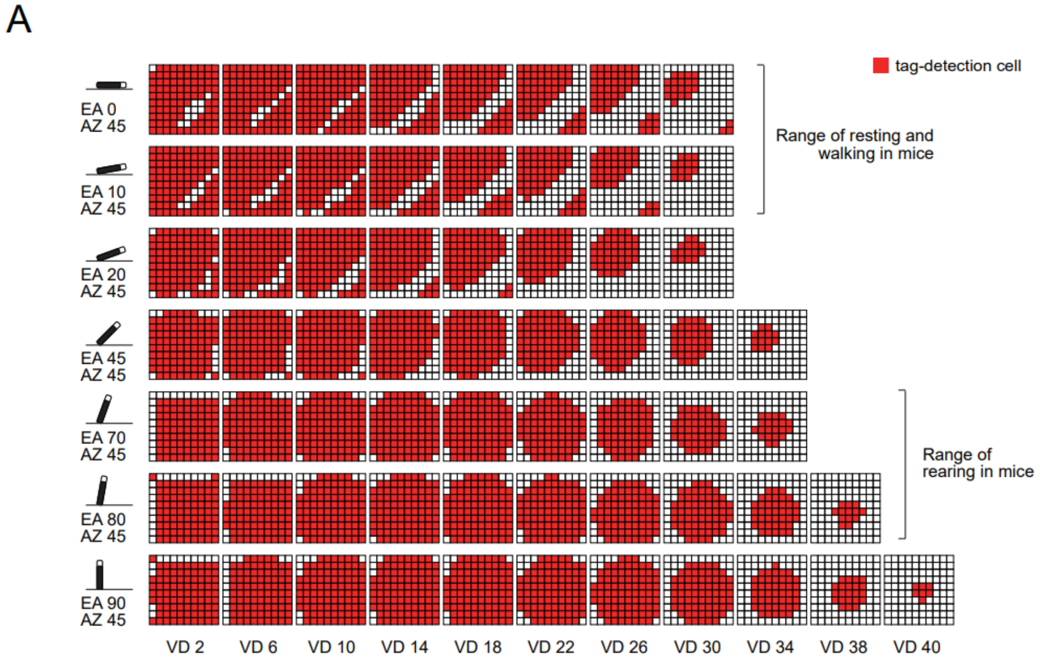


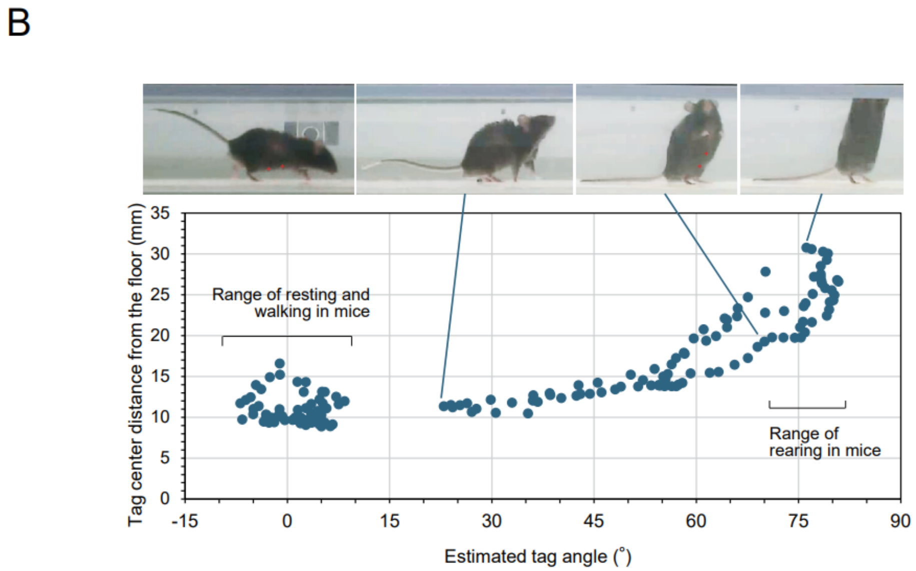


**Supplementary Figures for the Tag Detection Range.** (A) Tag detection performance across spatial configurations. Detection results are shown for all elevation angles (EA 0–90°) and vertical distances (VD 2–40 mm) under an azimuth angle of 45°. (B) Approximate position of the implanted tag in freely moving mice within the cage. Tag orientation and distance from the cage floor were estimated from video recordings of mice in different postures (walking, crouching, and standing) using MotoRater (TSE Systems, Germany). The distance from the floor (mm) was measured relative to the center of the estimated tag position. Tag angles for each posture were determined based on the contour of the ventral body surface.

**Supplementary Figure 2.**

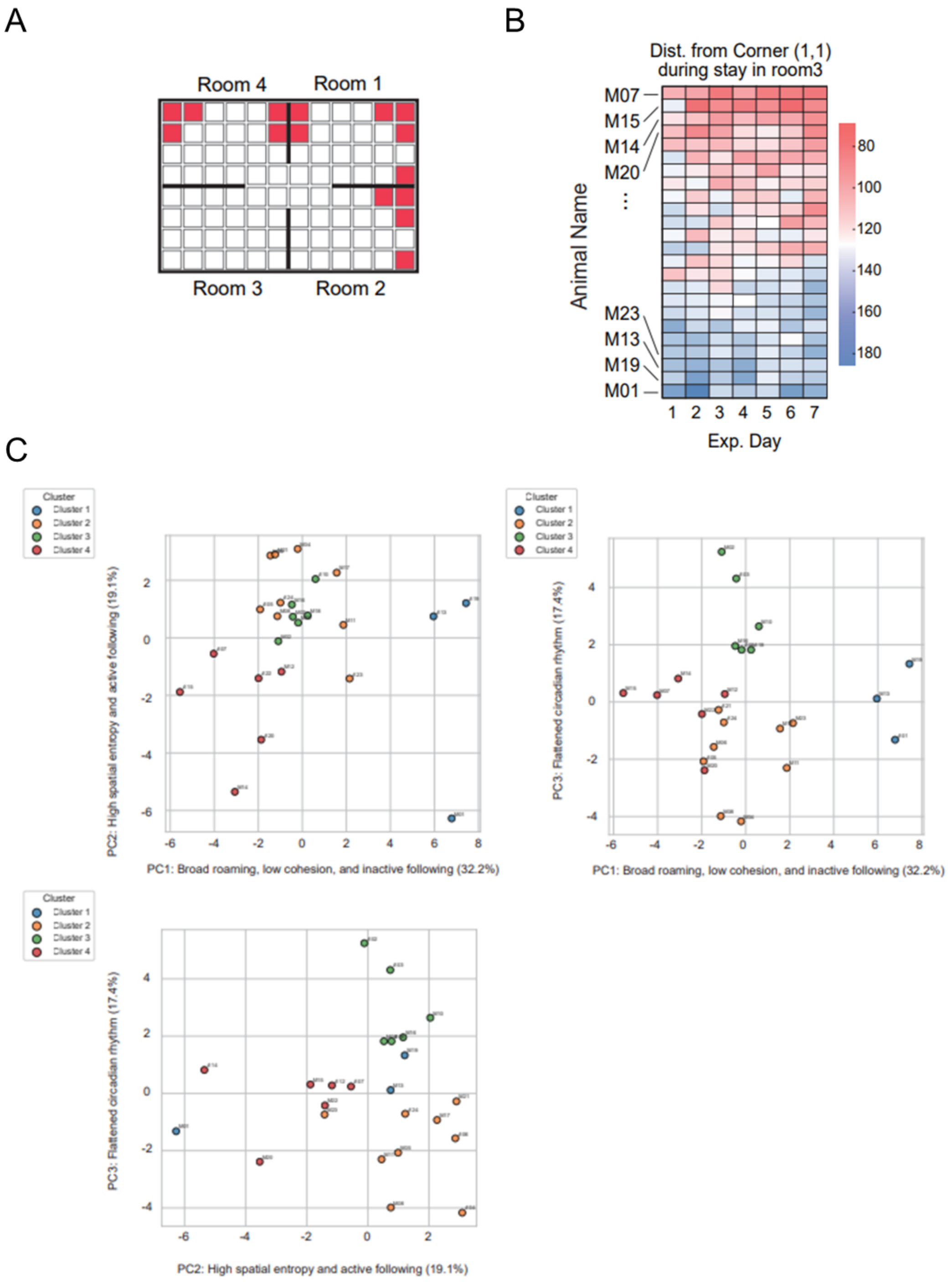


**Supplementary Figures for the Mice Experiment.** (A) Distribution of urine observed on the final day of the experiment (Day 7). The schematic highlights approximate urine locations in red. (B) Day-by-day changes in Distance from Corner (DFC) for each individual across the experimental period. (C) PCA biplot.

**Supplementary Figure 3.**

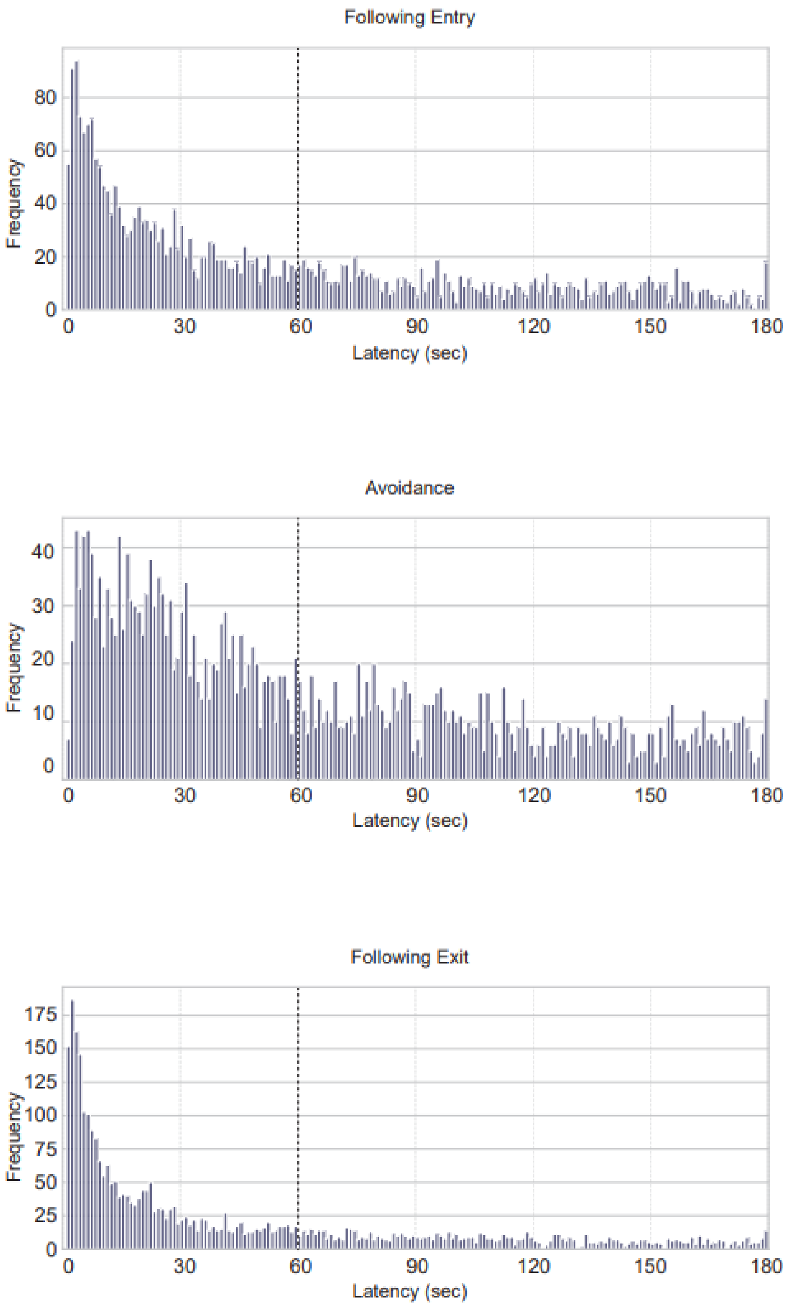


**Supplementary Figure for the Marmosets Experiment.** Top panel: Histogram of latencies for Following Entry (the time elapsed from when individual A enters Nest X until individual B enters the same Nest X), generated from all pair data. Middle panel: Histogram of latencies for Avoidance (the time elapsed from when individual B enters Nest X, where individual A is already present, until individual A exits), generated from all pair data. Bottom panel: Histogram of latencies for Following Exit (the time elapsed from when individual A exits Nest X until individual B exits the same Nest X), generated from all pair data.

**Supplementary Figure 4.**

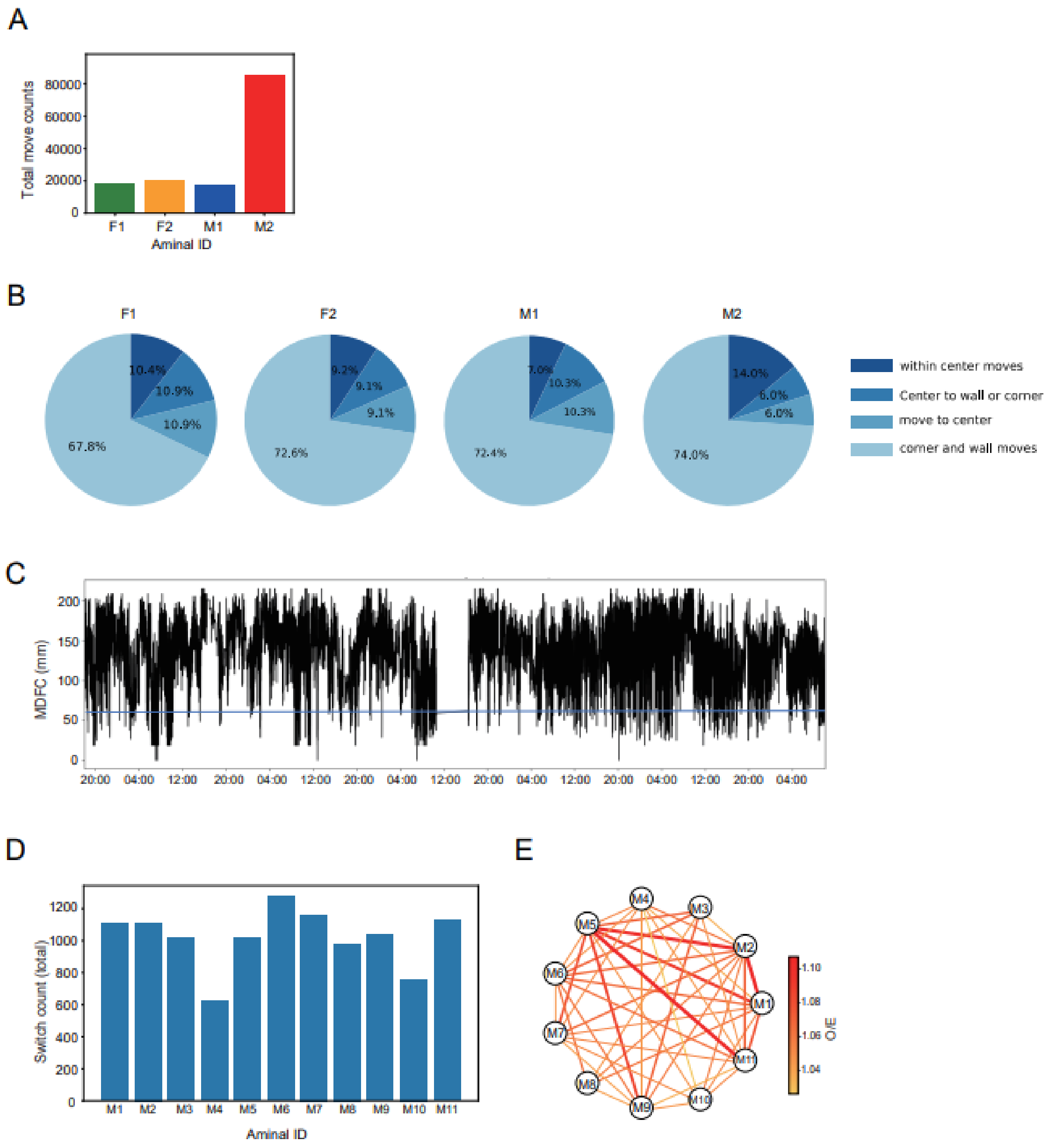


**Supplementary Figures for the Aquatic Environment Experiment.** (A) Total movement counts for each individual newt across the entire experimental period. (B) Movement categories. (C) Time series of group cohesion at 1-Hz resolution (1-s intervals), quantified as the mean distance from centroid (MDFC). The y-axis indicates MDFC (mm). They were fed around 18:00. (D) Total trafficking (compartment switching) counts per fish over the recording period. (E) Pairwise co-presence network. Undirected network constructed from pairwise same-compartment O/E ratios. Edges represent significant pairwise co-presence relationships (permutation test, $p < 0.05$).

**Supplementary Information 1. Supplementary Methods for Evaluation of Tag Read Range and Processing Speeds**

Because the coil of the cylindrical tag is wound around a ferrite core with high magnetic permeability, the magnetic flux is guided and concentrated within the coil even when its direction does not perfectly align with the longitudinal axis of the tag, enabling stable tag detection. Due to this effect, accurate estimation of the detection range by simple simulation is difficult; therefore, all evaluations in this study were performed experimentally.

For tag placement, seven elevation angles (0°, 10°, 20°, 45°, 70°, 80°, and 90°) and two azimuth angles (0° and 45°) were tested. The vertical distance from the antenna surface was set to 11 levels ranging from 2 to 40 mm (2, 6, 10, 14, 18, 22, 26, 30, 34, 38, and 40 mm). The horizontal position on the antenna surface was defined by dividing a 50 × 50 mm area into a 10 × 10 grid (100 cells). For all combinations of these conditions, whether the tag ID could be successfully read was recorded.

For the measurements, a 2-mm-thick transparent acrylic plate with a 5-mm grid printed on a transparent film was placed on the tile-type antenna to allow visualization of grid positions. A tag with a fixed elevation angle (using resin) was placed on the grid and moved horizontally while maintaining a constant azimuth angle. Tag readability was evaluated when the bottom end of the tag was positioned at the center of each grid cell, and the percentage of successful reads to total cells was defined as the Detection Area Coverage (%). The vertical distance was adjusted by stacking the 2-mm-thick acrylic plates, and Detection Area Coverage (%) was recorded for each condition.

The same procedure was applied to the ring-type antenna, except that cells corresponding to regions outside the circular coil were excluded from the analysis. In addition, tag reading distances in air, freshwater (hardness: 60–80 mg/L), and a 3.5% NaCl solution (Wako, Japan) were compared using the ring-type antennas. A ring-type antenna was lowered toward a tag fixed vertically ~2 cm above the bottom of a 3-L beaker. For liquid conditions, the antenna's magnetic field was thought to be fully submerged. Three independent antennas were tested.

Next, the polling cycle time was measured for 24 tile-type antennas in the 2D module and 24 ring-type antennas in the FLEX module. A single tag was placed at the center of each antenna (elevation angle 90°, vertical distance 0 mm), and the number of tags was gradually increased while measuring the polling cycle time. To evaluate the effect of anti-collision processing, multiple tags (up to 13) were placed on a single antenna, and the polling cycle time was measured for each condition. From the measured polling cycle time, the polling time per antenna and the tag read time per tag were calculated

**Supplementary Information 2. Supplementary information regarding the common marmoset diet**

During weekdays, the marmosets were fed solid food (CMS-1M, CLEA Japan Co., Ltd., Tokyo, Japan) soaked in room-temperature water for 5 minutes. Twice a week, the diet was supplemented with either sweet potato and powdered milk (Pig Liquid Natural for piglets, Scientific Feed Laboratory Co., Ltd., Tokyo, Japan) or honey water (prepared by dissolving 2000 g of honey, 50 g of vitamin D, and 50 g of vitamin C in water to a total volume of 2000 mL).

Snacks such as boiled rice crackers (Osaka Maeda Seika Co., Ltd., Japan) and castella cakes (Bourbon Co., Ltd., Japan) were provided biweekly during handling for weight measurement. On weekends, the animals received soaked solid food on one day and unsoaked solid food on the other. As needed, the diet was further supplemented with Biofermin (Biosuri for animals, Toa Pharmaceutical Co., Ltd., Japan; 4 g), high-protein milk (Meiji Co., Ltd., Japan; 4 mL), high-protein milk powder (Tube Diet High-Protein, Morinaga Sanworld Co., Ltd., Japan; 1.7 g), or Recovery Gel (DietGel® Recovery, ClearH2O, USA; 56 g per cup).

### Supplementary Information 3. Observational report by the marmoset caretakers

Period corresponding to behavioral data collection: April 18, 2023 – June 6, 2023

#### (A) Impressions of Behavior Toward Caretakers:

F1: Vigilant
F2: Friendly
M1: Cautious
M2: —
M3: Aggressive
M4: Vigilant

#### (B) Observational Impressions of Inter-individual Relationships During the Data Collection Period

**F1:** Good relationship with all other individuals (F1 dominant).

**F2:** Good relationship with F1 (F2 submissive), M1 (F2 submissive), M4 (F2 dominant). Poor relationship with M2 and M3.

**M1:** Good relationship with all other individuals (M1 dominant, except for F1, who is of equal status).

**M2:** Good relationship with F1 (M2 shows affiliative/dependent behavior), M1 (M2 submissive), M4 (M2 dominant). Poor relationship with M3 (M2 submissive), F2

**M3:** Good relationship with F1 (M3 affiliative), M1 (M3 rather subordinate at the beginning), M4 (M3 dominant). Poor relationship with M2 and F2 (M3 dominant)

**M4:** Good relationship with all other individuals (M4 submissive)

#### (C) Notes on Social Relationships and Animal Room Conditions (Including Pre- and Post-Experiment)

Until late January 2023, F1 accepted both M2 and M3 (embracement, mating).

**January 18, 2023**

M3 mated with F1; from this point until approximately May 2024, M3 and F1 formed a pair.

**January 23, 2023**

M2 attempted to mate with F1, but the attempt did not succeed because M3 interfered.

**February 24, 2023**

Food had been depleted and the group appeared hungry and irritated.
Feeding order observation: Observed order: M1 → F1 → M2 → M3 → F2.

**March 23, 2023**

M1 and F1 fought; hereafter, F1 came to act dominant.

**April 3, 2023**

Food ran out, F1 became irritated and threatened other individuals.
Feeding order observation: M1 ≈ F1 → (M2, M3, F2) → M4.
M1 was at the top, and F1 also competed strongly: F1 tried to eat first along with M1 and M1 scolded/reprimanded F1. However, F1 continued eating as if nothing had happened. M1 was strict toward M2, M3, and F2. M4 did not try to approach and stayed at the back.

**April 14, 2023**

M3 appeared to remain close to F1 and followed F1 constantly. F1 was presumed pregnant around this time (retrospective estimation).

**April 17, 2023**

M2 attempted mating with F1 but was noticed by M3 and interrupted, so the attempt ended unsuccessfully.

**April 18, 2023**

Room expanded. New individuals were also introduced (without chips).
**Initially, the housing area was divided into two compartments, with all tagged animals housed in one of the compartments. On the first day of data acquisition, the partition was removed to form a single contiguous space for behavioral tracking.*
M1 and F1 each threatened the newcomers, apparently to "teach"/establish hierarchy.
M2, M3, F2, and M4 did not show strong interest in the newcomers; they entered the new space and explored/played.
The newcomers (without tags) seemed afraid of individuals roughly up to F1, M1, M2, and M3. They showed submissive calls or hid in gaps/crevices. The newcomers did not appear to fear F2 or M4.

**April 27, 2023**

M3 followed F1 around. M2 continuously produced submissive vocalizations toward M3 and toward F1 (submissive, infant-like vocalizations were a typical vocal pattern observed as a sign of submission).

**April 28, 2023**

M2 repeatedly emitted submissive calls toward M3; M3 responded with threat vocalizations (video available). Previously, M2 and M3 often stayed together.

**May 1, 2023**

M2 produced submissive calls toward M3; M3 responded with threat vocalizations.

**May 2–May 9, 2023 (Holidays)**

Food and water were not replaced every day during the holiday period.

**May 9, 2023**

M2 produced submissive calls toward M3; M3 appeared angry/irritated in response.

**May 17, 2023**

The same pattern persisted: M2 continued submissive calling toward M3.

**May 18, 2023**

M2 produced submissive calls toward M3. M3 chased M2 while threatening; M2 fled.
This sequence (calling → chasing/threatening → fleeing) was repeated multiple times.

**May 19, 2023**

The same scene was observed again: M2 vocalizing toward M3, M3 threatening and chasing, M2 fleeing.

**May 22, 2023**

Box positions were changed (No. 5 & 6, and No. 15 & 16).

**May 24, 2023**

M3 repeatedly bit M2. M3 chased M2 persistently. Other individuals did not actively join or intervene, but appeared restless; the overall atmosphere was noisy/agitated.
M2 hid in narrow spaces (e.g., between the structure and wall) or behind logs.

**June 7, 2023**

M2 was bitten on the back by M3 and they were wrestling; the caretaker separated them.
After separation, M2 stayed quietly in the corner of the room, watching M3 cautiously.

**June 8, 2023**

As usual, M3 threatened; M2 remained vigilant and fled.

**June 15, 2023**

M2 provoked M3 using vocalizations; M3 chased M2. This did not escalate into a full wrestling fight, but the provoking/chasing sequence was observed frequently throughout the day.

**June 23, 2023**

A fight occurred between M2 and M3; M3 had a minor wound on the forehead. Even after being attacked, M2 quickly went back to approach M3 again.

**June 26, 2023**

M2 had a wound on the forehead (possibly due to fighting with M3).
M2 showed a decrease in body weight (366 g → 324 g). F1 showed a tendency toward weight gain.
*Note:* The repeated pattern was that M2 provoked, M3 became angry, and M2 fled; however, M2 did not appear to be trying to hide or escape permanently.

**August 7, 2023**

Food was almost empty; the individuals appeared very hungry (food had been replaced over the weekend).
F1 became angry if its feeding was disturbed and produced threat vocalizations; F1 was the dominant individual. Individuals other than F1 crowded around the three feeding sites and ate simultaneously. Unlike earlier periods, there was no longer an obvious pattern of allowing smaller individuals (like M4) to feed first. When M2 was eating, M3 came to interfere.

**August 10, 2023**

M2 appeared isolated and was vigilant toward M3. M2–M3 chasing continued. Previously, M2 had sometimes initiated interactions toward M3 while giving submissive calls, but recently the pattern was mainly initiated by M3. M3 approached the isolated M2, sat at a certain distance while asserting its presence, and then threat vocalizations and chasing began. M2 appeared highly wary of M3.

**August 15, 2023 (Holiday period)**

Food was empty; the group appeared hungry. M3 was seen chasing and threatening M2. F1 and M3 threatened others around the feeding area.
Because the group was restless/noisy overall, the caretaker entered and placed small amounts of dried food in multiple locations; the group ate and then calmed.

**August 17, 2023**

Food was empty; the group appeared hungry.
F1 initially monopolized feeding site A (Feeder 3,4) and ate first. maller individuals gathered at the site B(Feeder 1,2). M2 foraged for food that had dropped from the A/B feeding sites instead of feeding directly from the feeder boxes.

**August 28, 2023**

F1 threatened M1 (when feeding was interrupted).
While considering whether to install an additional wooden beam/log, all individuals showed interest at once except for M2.

**September 5, 2023**

F1 gave birth.

### Supplementary Information 4. Safety evaluation of HF RFID tag implantation

Mice were implanted at one month of age; 22 of 24 survived to 25 months, at which point tissue samples were collected, with no implant-related abnormalities detected upon inspection. All marmosets remain alive 3.5 years post-implantation as of this writing, with no abnormalities at the implantation site or general health. For newts and tilapia, two individuals of each species that were maintained for long-term observation remain alive for over 3 years after implantation, with no externally observable abnormalities.
